# Softening in two-component lipid mixtures by spontaneous curvature variance

**DOI:** 10.1101/2023.12.12.571323

**Authors:** Amirali Hossein, Andrew H. Beaven, Kayla Sapp, Alexander J. Sodt

## Abstract

The bending modulus of a lipid bilayer quantifies its mechanical resistance to curvature. It is typically understood in terms of *thickness*, e.g., thicker bilayers are stiffer. Here, we describe an additional and powerful molecular determinant of stiffness — the variance in the distribution of curvature sensitivity of lipids and lipid conformations. Zwitterionic choline and ethanolamine headgroups of glycerophospholipids dynamically explore inter- and intra-species interactions, leading to transient clustering. We demonstrate that these clusters couple strongly to negative curvature, exciting undulatory membrane modes and reducing the apparent bending modulus. Three forcefields (Martini 2, Martini 3, and all-atom CHARMM C36) each show the effect to a different extent, with the coarse-grained Martini models showing the most clustering and thus the most softening. The theory is a guide to understanding the stiffness of biological membranes with their complex composition, as well as how choices of forcefield parameterization are translated into mechanical stiffness.

## 1. INTRODUCTION

The mechanical properties of membranes are central to myriad vital processes in cells [1–3]. In contrast to single-component membranes typically used to deduce mechanics, biological membranes are comprised of hundreds of different lipid species [4], as well as proteins. Understanding biological membranes thus requires describing how mechanics changes in mixtures of many lipids with widely varying mechanical influence.

The spontaneous curvature (*c*0), bending modulus (*κ*), and Gaussian curvature modulus 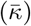 together parameterize the quadratic model describing the free energy cost of deviations of curvature from its preferred value [5, 6]. These constants can be determined from a variety of experimental and simulation-based methods. Common experimental methods for determining *κ* include optical analysis of shape fluctuations [7] and neutron spin echo spectroscopy [8]. With simulations, *κ* can be calculated *in silico* via thermal fluctuations of flat membranes [9], as well as equilibrium force methods such as buckling [10].Experimental values for *c*0 are typically found by analyzing membranes in inverse hexagonal phase [11] or from nuclear magnetic resonance spectroscopy studies of interleaflet redistribution of lipids in small unilamellar vesicles (SUVs) [12, 13]. In simulations, *c*0 is usually determined from the first moment of the lateral pressure profile [14, 15], or recently from free energy perturbation methods [16].

Given the complex composition of biological membranes, the relation between material parameters of lipid mixtures and those of their constituent components is of great interest. A reasonable assumption would be to expect mechanical parameters of multi-component membranes to be weighted averages of those corresponding to pure membranes of each of the components [17]; however, simulation studies show in many cases of biological importance this expectation of additivity of mechanical properties (e.g. spontaneous curvature) does not hold true [18]. Furthermore, it has been pointed out that a purely shape-based description of membrane elements fails to take into account the effect of the environment surrounding a molecule on its curvature preference [19], such as hydrogen bonding between adjacent lipids. Cholesterol, a ubiquitous part of many biological membranes, appears to have a particularly non-trivial influence on mechanical properties of membranes; its addition to pure bilayers of different phospholipids has been observed to increase or not alter the bending modulus of the bilayers, depending on the level of unsaturation of the tail acyl chains [20].

The underlying mechanism driving the phenomenon is the lateral diffusion of curvature-sensitive membrane species at time-scales differing from the characteristic relaxation times of undulation modes, leading to enhancement of their amplitudes and a softer apparent bending rigidity observable at longer time-scales. This coupling between local curvature and composition which, to the best of our knowledge, has first been described in literature by Leibler [21] (and subsequently addressed further theoretically [22, 23] and observed experimentally [24, 25]), is here treated for the case of multicomponent bilayers and its intricacies is investigated using simulations of binary mixtures.

The softening of complex mixtures would most clearly be indicated by a simple model of a system well-characterized experimentally at long length-scales: a mixture of phosphatidylethanolamine (PE) and phosphatidylcholine (PC). Here, the spontaneous curvature difference between coarse-grained PE and PC (Martini versions 2 and 3, abbreviated M2 and M3) is computed using three complementary and independent methods: the literature standard lateral pressure profile method [14, 15], thermodynamic integration (TI) of lipids into the inverse hexagonal phase [16], and the spatial extent (SE) methodology tracking the dynamic redistribution of lipids on fluctuating membranes [26, 27].

Curvature redistribution of Martini PE and PC lipids in mixtures between 0 and 100% PE indicates the expected spontaneous curvature differs between the two lipids. PE bilayers are found to be substantially softer. The spontaneous curvature of PE and PC lipids is further broken down by the number of attractive headgroup-headgroup interactions a lipid experiences. PE forms more higher-order clusters of attractive interactions — these lipids have very negative spontaneous curvature. A multi-component softening theory and analysis indicates that the presence of these higher order clusters should lead to substantial softening of the PE-rich bilayers. Complementary analysis of all-atom simulations indicate that all-atom PE does not form such clusters to nearly the same extent, and that all-atom PE bilayers are not as soft, relative to all-atom PC. The consequences of this observation are discussed both in terms of designing coarse-grained forcefields as well as understanding the impact of nanoscale heterogeneity on mechanics.

## II. METHODS

### A. System build information

#### 1. All-atom

All-atom planar bilayers were built at 0:100, 50:50, and 100:0 mol% DOPC:DOPE with 200 lipids/leaflet using the CHARMM-GUI *Membrane Builder* module [28, 29]. Each membrane composition was built with three independent replicas. The bilayers were solvated with 50 H2O/lipid and no ions.

#### 2. Coarse-grained for thermodynamic integration (TI)

All coarse-grained systems were built using M2 [30, 31] and M3 [32] lipid representations that reduce DOPC and DOPE to 12-bead representations. Within a lipid model (e.g., either M2 or M3), the DOPC and DOPE lipids differ by only the head group bead. Lipid inverse hexagonal phase (HII) systems were built [16, 33] at 0:100, 50:50, and 100:0 mol% DOPC:DOPE with initial radii of 30, 35, 40, 65, and 90 Å. The radius is defined by the `C1A` bead, which is the approximate location of the pivotal plane (the site at which lipids bend with constant area) [16, 34, 35]. All HII systems had 400 total lipids except for the HII system with an initial radius of 90 Å that contained 500 total lipids. Hexadecane (in Martini, four straight-chain saturated acyl-chain interaction sites) was placed in interstitial spaces to relax the hexagonal packing geometry [36, 37]. These systems have 2635, 3179, 3717, 6347, and 11108 water beads, respectively, and no ions. Planar bilayers were built using modified CHARMM-GUI *Martini Maker* scripts [28, 38] with 200 lipids/leaflet at 0:100, 50:50, and 100:0 mol% DOPC:DOPE. Hexadecane was also placed in the center of planar bilayers using PACKMOL [39] so that the bilayer is an applicable reference state for HII simulations. All planar systems had *∼*10 water beads/lipid and no ions. All systems (HII and planar) were built in triplicate.

### 3. Coarse-grained for fluctuation analysis

Large coarse-grained bilayers (1000 lipids/leaflet) were built at 0:100, 10:90, 20:80, …, 100:0 mol% DOPC:DOPE to interpret the undulation spectrum and dynamic redistribution of lipids. The CHARMM-GUI *Martini Maker* [40, 41] module was used to build these lipid bilayers with M2 [30, 31] and M3 force fields [32]. The simulations contained approximately 10 CG water beads per lipid and no ions.

### B. Simulation

#### 1. All-atom for fluctuation analysis

After a brief minimization and equilibration using NAMD [42, 43] with the CHARMM all-atom force field [44], the systems were converted to Amber format [45, 46] using ParmEd. These simulations were used to calculate fluctuation data. Each replica was simulated 2 µs using the Amber22 version of `pmemd.cuda` [47–49] and the CHARMM all-atom C36 force field[44]. Semi-isotropic pressure (1 bar) was maintained by a Monte Carlo barostat [50, 51], and constant temperature (310.15 K) was maintained by Langevin dynamics with a 1 ps^-1^ damping coefficient. Covalent bonds involving hydrogen were constrained using the SHAKE and SETTLE algorithms [52, 53]. Non-bonded forces were switched off between 10–12 Å, and long-range electrostatics were calculated by PME [54, 55].

#### 2. Coarse-grained for thermodynamic integration (TI)

All systems were minimized, equilibrated, and simulated using GROMACS 2019.3 [56–58]. The temperature was maintained at 310.15 K by velocity rescaling [59] with a time constant of 1 ps. Semi-isotropic pressure of 1 bar was maintained by a Parrinello-Rahman coupling algorithm (compressibility = 3 ×10^−4^ bar^−1^, pressure coupling constant = 12 ps) [60, 61]. A 20 fs time step was used, and the dielectric constant was set to 15. Each system was simulated 3 µs with both the M2 [30, 31] and M3 [32] force fields. The independent system configurations from 1, 2, and 3 µs were used as seeds to start the TI simulations.

For each of these three checkpoints, 11 *λ* windows were initiated (*λ* = 0.0, 0.1, …, 1.0) and simulated for 100 ns with a 20 fs time step (TI run parameters used were scalpha = 0.5, sc-power = 1, and sc-r-power = 6). The average was obtained from these three blocks, leaving off the first 5 ns of all simulations to allow configurational equilibration. After the simulations, the free energy derivative at each *λ* (⟨*U* ^*′*^⟩(*λ*)) was integrated using Bennett’s acceptance ratio (BAR; `gmx bar`) algorithm, yielding Δ*F*A*→*B.

#### 3. Coarse-grained for 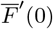

The 0:100, 50:50, and 100:0 mol% DOPC:DOPE planar bilayers containing 200 lipids/leaflet were minimized, equilibrated, and simulated using NAMD. Again, the M2 [30, 31] and M3 [32] force fields were tested. M2 force field parameters in NAMD format were taken from CHARMM-GUI [40, 41] and M3 force field parameters in NAMD format were made for this manuscript (available at: https://github.com/beavenah/martini3_namd). Specific to using the Martini force field in NAMD: cosAngles and martiniSwitching options were on (switch distance of 9 Å and a cutoff of 12 Å), PME was off, a dielectric of 15 was used, and a 20 fs time step was used. Pressure was maintained at 1 atm by a Nosé-Hoover piston (piston period of 2000 fs and decay of 1000 fs) and a temperature of 310.15 K was maintained by a Langevin piston. All systems were simulated 600 ns.

#### 4. Coarse-grained for fluctuation analysis

Larger (1000 lipids per leaflet) bilayers were simulated for 6 µs using the M2 [30, 31] and M3 [32] force fields using the GROMACS 2019.3 package [56–58]. The simulation parameters (e.g., temperature, pressure, etc.) were the same as those listed in the TI simulation subsection.

### C. Thermodynamic integration (TI)

TI computes the change in free energy (Δ*F*) by alchemically mutating “species A” into “species B” through intermediate parameter space. The method uses a *λ* parameter that shifts the system’s potential energy (*U*A→*U*B): *U* (*λ*) = *U*A + *λ*(*U*B−*U*A), for example, between DOPC and DOPE. For the DOPC↔DOPE transformation in the Martini force field, only intermolecular interactions are changed (i.e., the bonded and angle parameters are constant with *λ*). For the M2 and M3 force fields, the head group bead Q0 (M3: Q1) of DOPC is mutated to Qd (M3: Q4p) for DOPE.

For each membrane composition and phase, TI was run to obtain 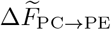 and 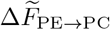 (where the tilde represents a per-lipid quantity). These 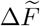 quantities include curvature-dependent and curvature-independent contributions to the free energy. We assume that bending a leaflet from a planar reference state only changes the curvature contribution to the free energy, therefore:

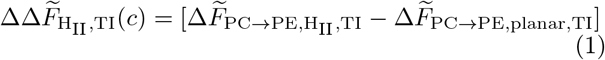

can be modeled by a Helfrich-Canham Hamiltonian that considers only curvature.

The per-lipid curvature free energy for a lipid X in phase Y is given by:

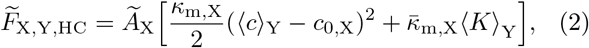

where lipid X’s per-lipid area, leaflet bending modulus, intrinsic curvature, and Gaussian curvature modulus are Ã_X_, *κ*_m_,X, *c*_0_,X, and 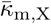, respectively. The phase’s mean and Gaussian curvature at the neutral surface are ⟨*c*⟩Y and ⟨*K*⟩_Y_, respectively. The planar phase has zero ⟨*c*⟩ and ⟨*K*⟩ while the HII phase has zero ⟨*K*⟩.

As shown previously [16], assuming that *κ*m and 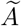 are equal for DOPC and DOPE (that is, that *κ* is homogeneous):

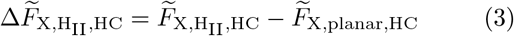

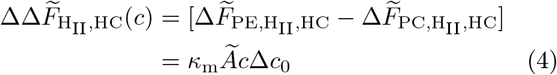

As an example, the DOPE is the final state mutated from the initial DOPC state. The planar phase is used as the reference state relative to the HII phase.

For each HII radius, there exists a calculated 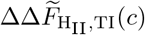 and a predicted 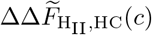. For 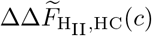, we calculated *κ*m from methods developed herein, Ã from the simulation box size, and *c* from the HII applicable simulation. The *κ*m and Ã are the average of the calculated DOPC and DOPE values. Calculating the HII radius requires finding the lipidic pivotal plane (where lipids bend at constant area). To do so, the Ã was calculated from the bilayer+alkane simulations to account for alkane-induced lipid area dilation. Then, the average area and radius of each lipid bead was calculated for each HII system [16, 35]. The lipidic location whose area matches the bilayer+alkane lipid area is the pivotal plane (e.g., Ref. [16]). Note that the radius need not correspond to a bead. In these simulations, the pivotal plane lies between the `GL2` and `C1A` beads, but typically much closer to the `C1A` bead.

The fit between 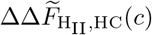 and 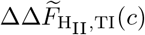 was obtained by a least-squares fit with Δ*c*0 as the only parameter. The error in the fitted Δ*c*0 parameter was calculated by fits to Monte Carlo generated data sets that sample the standard deviation of Δ*c*0 based on the statistical error of the TI data.

### D. Free energy derivative with respect to curvature 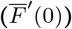

For each bilayer+alkane system, the free energy derivative with respect to curvature 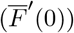 was also calculated using a patched version of NAMD 2.12 [62].

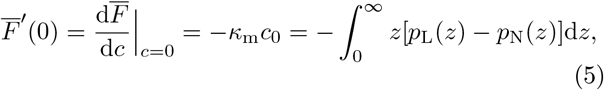

where *c* is the leaflet curvature at the pivotal plane, *z* is the vertical distance from the bilayer midplane, and *p*L and *p*N are the lateral and normal components of the pressure tensor, respectively. By convention, a positive 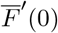 indicates a leaflet that would bend toward its head groups if unconstrained. The 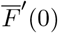 magnitude indicates curvature stress (compare to Eq. 4).

### E. Calculating the bending rigidity (*κ*b) from membrane fluctuations

The bending modulus of the simulated systems in this work are calculated using two different methods, both of which follow from Helfrich-Canham (HC) continuum elastic theory of membranes [5, 6].

In the first approach, we calculate the height fluctuation spectrum of bilayers from simulations and find bending rigidity modulus *κ*b by fitting the data for amplitudes of fluctuation modes to the theoretical expectation from equipartition theorem:

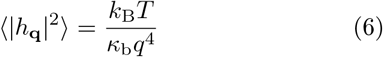

Here *h***q** is the amplitude of the Fourier mode **q** of the surface height, which we calculate by discretizing the bilayer surface with lattice length of *L*≈15Å, and using the lipid atom corresponding to the neutral surface of the leaflet (discussed below) to determine surface height in each lattice point.

In the second method, we extract the bending rigidity modulus from the slope of the transverse curvature bias (TCB) [26]:

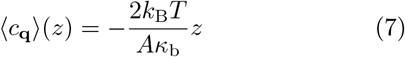

Here ⟨*c***_q_**⟩ (*z*) is the mean curvature calculated from mode **q** at distance *z* away from neutral surface of the leaflet along the surface normal. The reference atom (or “bead” in coarse-grained Martini model’s terminology) used to represent the leaflet neutral surface is determined by finding the surface parallel to the bilayer midplane where the sampled curvature of atoms is not biased toward positive or negative values.

We argue the TCB method is preferable to the height fluctuation method due to sensitivity of the latter to lipid tilt/protrusion fluctuations at higher *q* values [63]. The mean sampled curvature of the lipid species in the mixed systems is calculated to extract information about their intrinsic curvature preferences as revealed by the coupling between lateral distribution of the lipids and undulation modes of the bilayer surface.

### F. Membrane softening theory

#### 1. Diffusional and conformational softening

Two mechanisms are expected to potentially reduce the “apparent” bending rigidity of a membrane mixture [64]. The first mechanism arises from the coupling between lateral distribution of lipid species with different curvature preferences and fluctuations in local curvature; This is referred to as “diffusional softening”, and, for a membrane consisting of a small fraction *χ* of components with spontaneous curvature Δ*c*0 relative to the background lipids, is described by

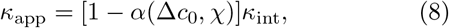

where *α*(Δ*c*0, χ) is a softening factor given by

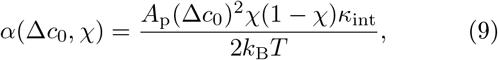

where *A*p is the area of the molecular membrane component.

A mathematically equivalent softening factor can also influence the apparent rigidity of membranes arising from formation of molecular complexes with a different curvature preference than what would be expected from a naive additive model, i.e., the area-weighted average of the components of the multi-molecular complex; now *χ* represent the fraction and Δ*c*0 the difference in intrinsic curvature of the curvature sensitive conformation.

#### 2. Multi-component softening

The lipids involved in the coarse-grained equivalent of a hydrogen bond (*pseudo* hydrogen bonds in M2 and M3 models) are determined by tagging the lipids with their head beads (`NH3` for DOPE and `NC3` for DOPC) closer than a threshold value of 6 Å to the phosphate (`PO4`) bead of an adjacent lipid. We group lipids with respect to their type and number of interactions (in which the considered lipids act as the head-group donor) and conduct a similar curvature-spectrum analysis for each subset of the lipids separately to distinguish between their relative curvature preferences.

For atomistic systems we use the distance between the N and P atoms to identify the inter-lipid interactions, with the same cut-off length as the CG simulations.

Applying the framework of 2-component diffusional softening to our PE/PC mixtures does not allow us to explain the difference in bending rigidity of the pure membranes. With this motivation, in order to be able to take into account the effect of all the subgroups identified based on inter-molecular interactions, we expand the softening theory to the case of many components. We find (Supplementary Materials) that for a system consisting of *n* lipid species with spontaneous curvature *c*0,*i* and composition fraction χ*i*, and intrinsic bending rigidity *κ*int:

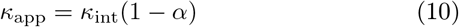

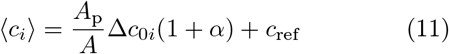

with

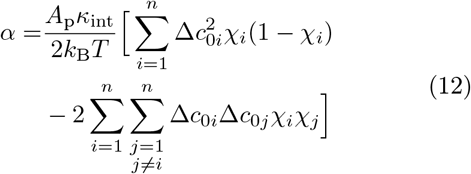

where here *c*ref is a reference curvature, i.e., the mean spontaneous curvature relative to which curvatures are measured in this method. Solving Eqs. 10 and 11 for *κ*_int_ and the set of {Δ*c*0,*i*} yields:

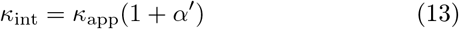

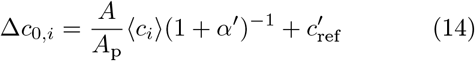

with

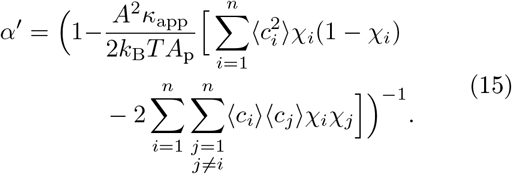

For the case of a distribution of spontaneous curvatures with variance *σ*^2^:

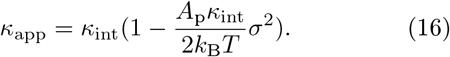

In the SE methodology, *c*ref is set to the mean, calculated as:

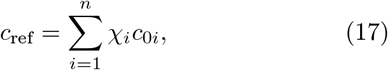

such that all observed and spontaneous curvatures are reported as Δ*c*s, relative to the reference.

## III. RESULTS AND DISCUSSION

### A. TI

The fit to TI data yields Δ*c*0 of –0.02456 ±0.00002 Å^−1^ (M2) and –0.02053 ± 0.00005 Å^−1^ (M3) with the minus indicating a more negative curvature preference for DOPE. This analysis uses a *κ*_m_ of 8.68 *k*_B_*T* for M2 (*κ*_m_,DOPC = 10.37 *k*_B_*T* and *κ*_m_,DOPE = 6.98 *k*_B_*T*) and a *κ*_m_ of 6.23 *k*_B_*T* for M3 (*κ*_m_,DOPC = 7.53 *k*_B_*T* and = 65.6 Å^2^). *κ*_m_,DOPE = 4.93 *k*_B_*T*). Additionally, an Ã of 66.5 Å^2^ for M2 (Ã DOPC = 68.4 Å^2^ and Ã DOPE = 64.6 Å^2^) and an Ã of 67.2 Å^2^ for M3 (Ã DOPC = 68.9 Å^2^ and Ã DOPE= 65.6 Å^2^).

These Δ*c*_0_ values compare well to previously published experimental and simulation data for DOPC/DOPE [35, 37, 65–68]. All-atom simulations of the planar and hexagonal phase using the CHARMM C36 force field yielded *c*0,DOPE = −0.038 Å^*−*1^ and *c*0,DOPC = −0.015 Å^*−*1^ (Δ*c*_0_ = –0.023 Å^−1^) [35]. Using only planar bilayer simulation data with the CHARMM C36 force field gives −0.027 and −0.007 Å^*−*1^ for DOPE and DOPC, respectively (Δ*c*_0_ = –0.020 Å^−1^) [65]. The values compare well to X-ray diffraction experiment results on the inverted hexagonal phase radii: –29.6 Å and –87.3 Å, for DOPE and DOPC, respectively (Δ*c*_0_ = –0.0223 Å^−1^) [37, 66, 67].

### B. Curvature spectrum

We look at the contribution of different undulation modes to total sampled curvature of the lipids (curvature spectrum) to determine the spatial extent of the lipids, i.e., their influence on the mechanical properties of the surrounding membrane. We observe a relatively local spatial extent for both lipids, with positive and negative values of spontaneous curvature relative to the background for DOPC and DOPE, respectively, as expected for M2, M3, and all-atom C36 simulations (Figs. 1, 2, and 3).

**Figure 1:**
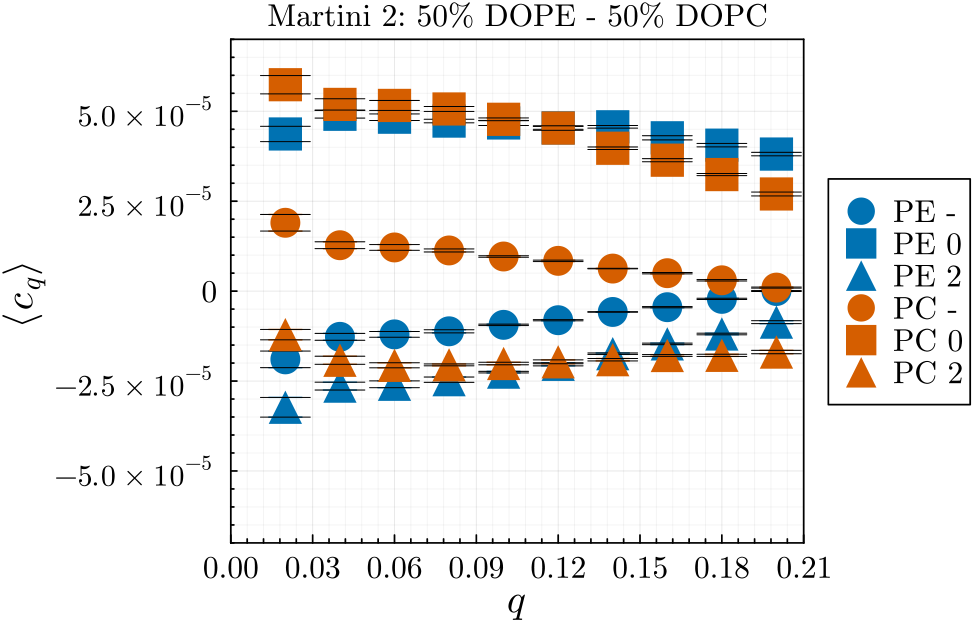
Sampled curvature spectrum of 1:1 DOPE and DOPC membrane in M2 (circles). The curvature spectrum for the subset of each type of lipids identified as having formed 0 (squares) or 2 (triangles) H-bonds is also shown.

**Figure 2:**
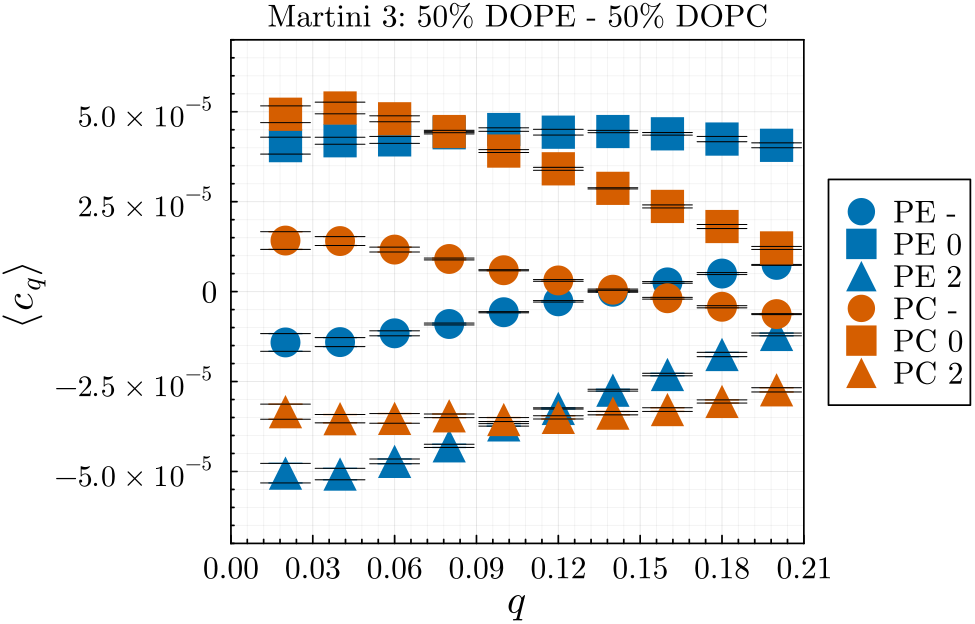
Sampled curvature spectrum of 1:1 DOPE and DOPC membrane in M3 (circles). The curvature spectrum for the subset of each type of lipids identified as having formed 0 (squares) or 2 (triangles) H-bonds is also shown.

**Figure 3:**
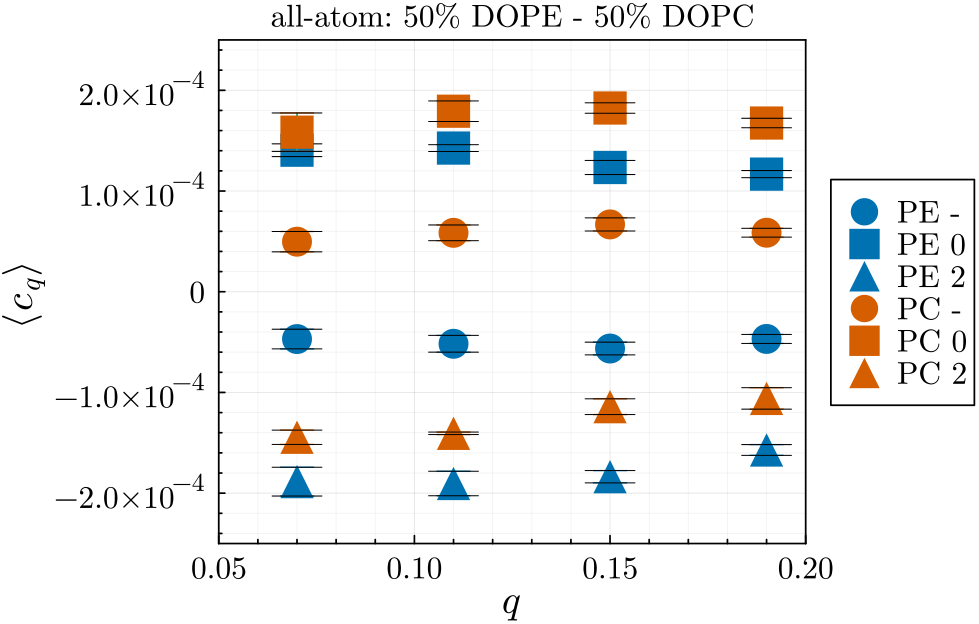
Sampled curvature spectrum of 1:1 DOPE and DOPC membrane in the all-atom C36 model (circles). The curvature spectrum for the subset of each type of lipids identified as having formed 0 (squares) or 2 (triangles) H-bonds is also shown.

However, looking at the curvature spectrum of the subspecies identified based on the number of inter-lipid interactions, we observe that lipids with higher number of pseudo hydrogen bonds (as defined for each model in Sec. II F 2) have a more negative curvature preference regardless of their type (PE/PC), while lipids with no interactions have a more positive curvature preference. We also observe that due to the different parameterization of the DOPE and DOPC head beads in the Martini model (both M2 and M3), the former has a greater tendency for forming such interactions, leading to a larger fraction of PE lipids being in the subgroup with two interactions, which in turn causes the overall more negative curvature preference of PE lipids as a whole group.

### C. 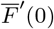

By way of Equation 5, Δ*c*_0_ can be calculated independently from the curvature spectrum. Using *κ*_m_ calculated herein, the values of Δ*c*_0_ are too large for both M2 and M3 (− 0.046±0.003 and − 0.034±0.003 Å^*−*1^, respectively). The excellent agreement of Δ*c*_0_ for TI and the curvature spectrum (see Fig. 7) suggests a discrepancy between the *local* Δ*c*_0_ and the *global* leaflet spontaneous curvature that begs further investigation.

### D. Apparent and intrinsic bending modulus

For M2 and the all-atom systems, the computed bending rigidities deviate from a linear sum of pure PC and pure PE membranes weighted by their mole fraction (Figs. 4 and 5). In particular, we observe a significantly softer apparent stiffness at 50% PE for M2, as expected due to the increased spontaneous curvature variance of a mixture. For M3, the spontaneous curvature variance is *less* for the 50%/50% mixture as computed from the tagged spontaneous curvature distribution.

**Figure 4:**
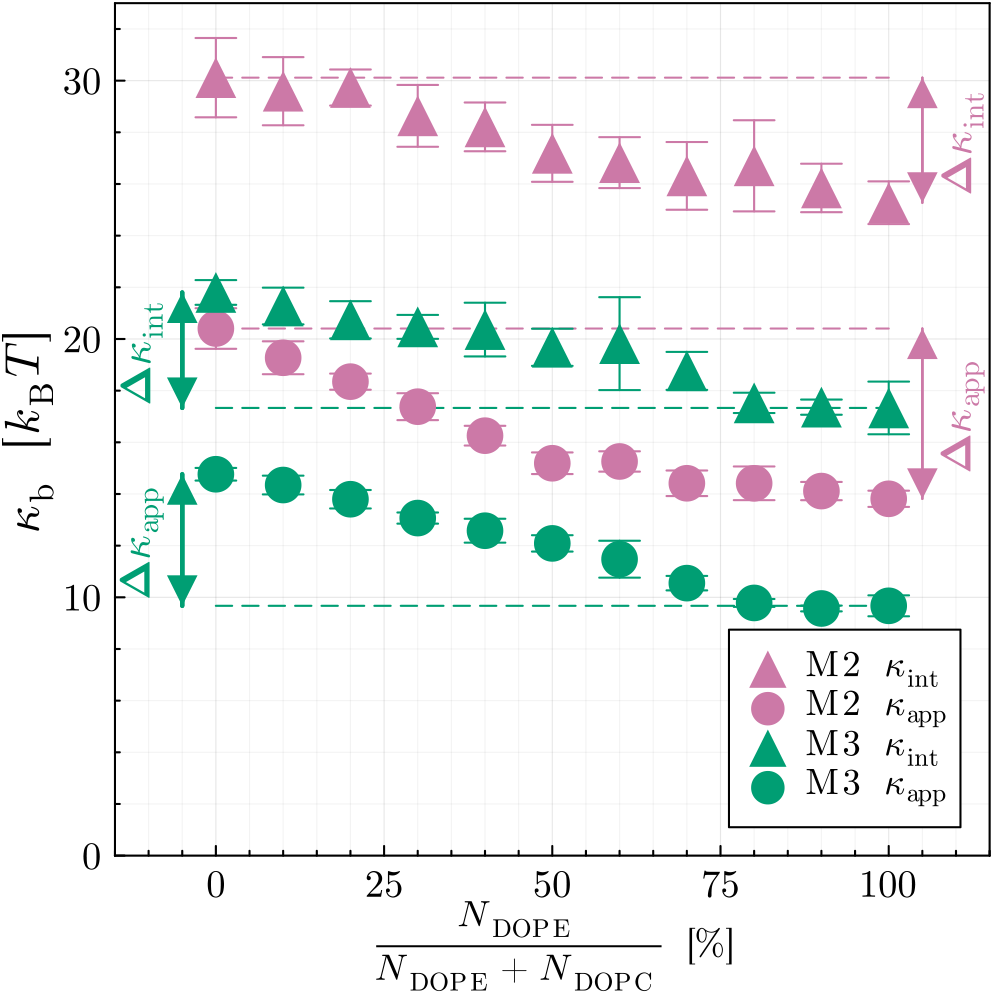
Apparent and intrinsic bending rigidity modulus for Martini DOPE/DOPC binary systems with varying compositions.

**Figure 5:**
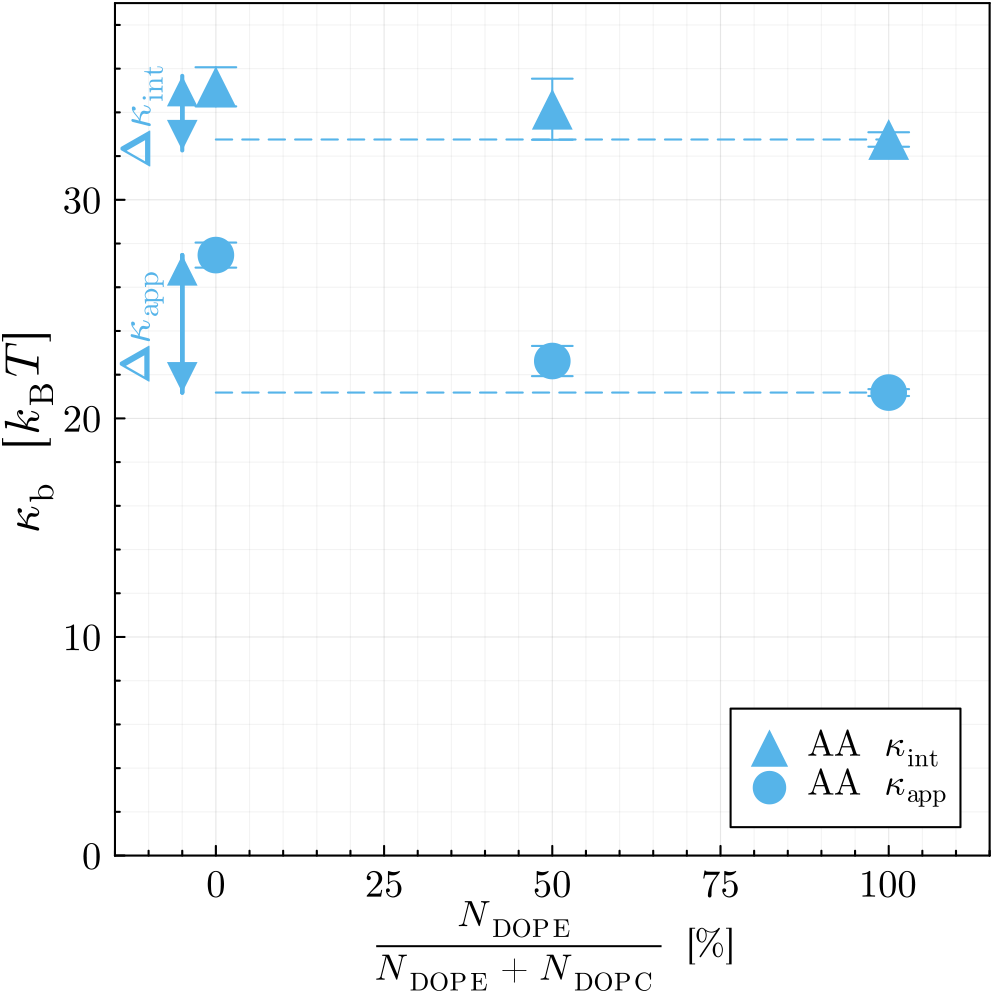
Apparent and intrinsic bending rigidity modulus for all-atom C36 DOPE/DOPC binary systems with varying compositions.

The calculated *κ*int are relatively constant with respect to composition. The softening with increased PE is not completely eliminated. However, our analysis focuses on a specific, discrete selection of subspecies that may not perfectly capture the difference between PE and PC. These results strongly underline the main idea behind the concepts of diffusional and conformational softening, which is that a diversity in curvature preference of membrane species can couple with the undulation modes of the membrane to lead to a softer apparent bending modulus.

## IV. CONCLUSIONS

In this work we have enriched an important mechanism by which lipid interactions and lipid clusters soften equilibrium measurements of the bending modulus, compared to the expectation from an idealized diffusion-static, noninteracting picture. As in Ref. [64] we refer to the former as the *apparent* modulus and the latter as the *intrinsic* modulus. The term “apparent” should not in any way diminish the value of this bending modulus — it is indeed the equilibrium bending modulus that is critical to most biological membrane reshaping.

First, our simulations show that, by accounting for the variance of the spontaneous curvature distribution, mixtures of lipids with differing curvature preference are indeed softer than area-weighted averages of their corresponding pure membranes, confirming theoretical predictions on the basis of the coupling between slow-relaxing diffusion modes and undulation modes. An exception was the M3 100% DOPE simulation, which had a larger spontaneous curvature variance than 50% DOPC/50% DOPE when head-group donating clusters were accounted for (Fig. 6). This indicates that, in this case, *conformational* softening dominated the effect of diffusional softening.

**Figure 6:**
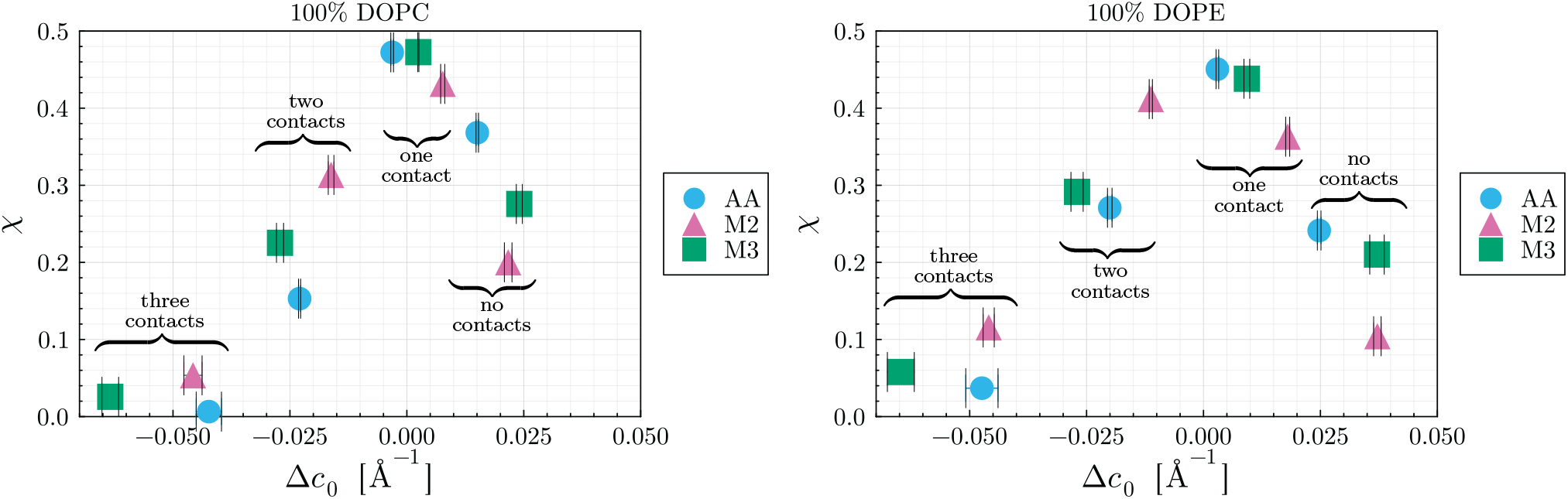
Fractional population of pseudo hydrogen-bonded configurations versus spontaneous curvature. For each simulation (M2, M3, all-atom) configurations are tagged donating three, two, one, or zero choline/ethanolamine to phosphate interactions (donating three interactions has the most negative Δ*c*_0_). The all-atom distribution is the most narrow, corresponding to a smaller softening constant, *α*.

**Figure 7:**
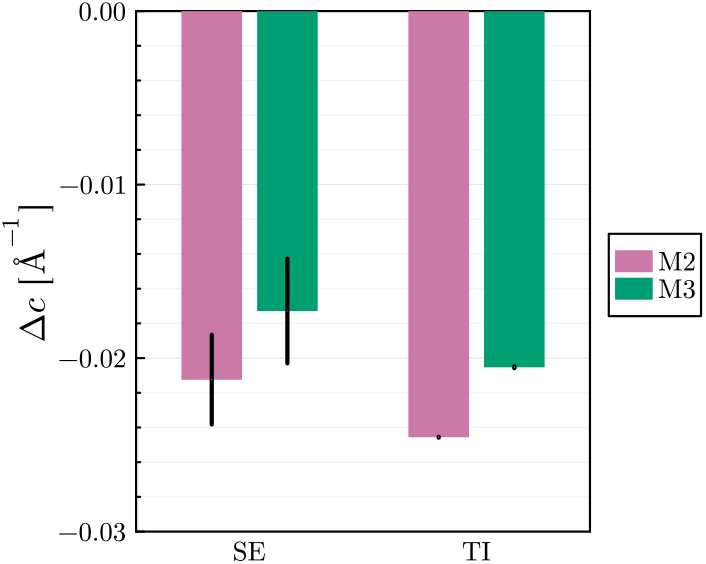
Spontaneous curvature difference between DOPE and DOPC lipids in Martini (M2 and M3) models, calculated using Thermodynamic Integration (TI) and Spatial Extent (SE) methods.

Second, we have measured the difference in spontaneous curvature of two lipid types using two alternative MD-based methods, and show that their results are in agreement, generating more confidence that each method can be used to reliably estimate membrane models curvature elastic parameters. Fig. 7 compares the calculated difference in spontaneous curvature of DOPE and DOPC lipids in M2 and M3 models calculated from TI and SE methods. While methods directly measuring Δ*c*_0_ are in excellent agreement, the lateral pressure profile method (see Table 1) yield spontaneous curvature estimates that were too high, suggesting a significant non-local impact of M2 and M3 lipids on curvature stress.

**Table 1:** 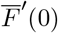 data for the Martini (M2 and M3) force fields with NAMD. Dividing 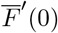 by −*κ*_m_ yields the leaflet spontaneous curvature.

| Force field (w/, w/o additive) | $\bar{F}'(0)_{100:0}$ [$k_B T/\text{\AA}$] | $\bar{F}'(0)_{50:50}$ [$k_B T/\text{\AA}$] | $\bar{F}'(0)_{0:100}$ [$k_B T/\text{\AA}$] |
| --- | --- | --- | --- |
| **M2 (+hexane)** | $0.4780 \pm 0.0052$ | $0.4082 \pm 0.0034$ | $0.3240 \pm 0.0041$ |
| **M2 (lipid-only)** | $0.4908 \pm 0.0039$ | $0.3604 \pm 0.0010$ | $0.2447 \pm 0.0018$ |
| **M3 (+hexane)** | $0.4074 \pm 0.0068$ | $0.3323 \pm 0.0029$ | $0.2809 \pm 0.0034$ |
| **M3 (lipid-only)** | $0.3956 \pm 0.0041$ | $0.3086 \pm 0.0034$ | $0.2393 \pm 0.0034$ |

Additionally, we have examined the role played by the transient curvature-sensitive clusters in determining the apparent bending rigidity of membranes, highlighting the importance of the ability to form such configurations in the softening effect. Specifically, we explain a significant portion of the difference in simulated apparent bending modulus of pure DOPE and DOPC membranes by dividing lipids into curvature-sensitive sub-groups based on inter-molecular interactions and using the multi-component softening theoretical framework to extract intrinsic quantities. At 100% DOPE for the M2, M3, and C36 models, *κ*_app_ is 68%, 66%, and 77%, while *κ*int is 84%, 80%, and 93% of the DOPC value, respectively. Note that the *absolute* gap between DOPE and DOPC *κ*_int_ is reduced by 26%, 12%, and 62%, for the M2, M3, and C36 models, respectively. This demonstrates how differences in chemical parameterization of the two lipids in three considered force fields leads to differences in molecular interactions that in turn alter the relative populations of multi-lipid complexes with distinct curvature preferences. Ref. [69] notes that “top-down” parameterized coarse-grain models (akin to Martini) may have problems being extended outside their range of parameterization, especially if they don’t reproduce key struc-tural details. Here, although qualitatively the behavior is similar, both DOPE and DOPC are overly clustered relative to the all-atom, leading to a softer bilayer. Note that this “intrinsic” *κ* attempts to remove the impact of the curvature sensitivity of the choline/ethanolamine-to-phosphate attractive interaction on *κ* — it is not a real mechanical constant and would only be of use if the clustering effect of PE/PC were to be explicitly modeled. Similar logic applies to an intrinsic constant modeling the bending modulus of a mixture of two or more components; it would only be of use if the lateral dynamics of the components were included explicitly. That is, if the lateral dynamics of a two-component mixture of curvature-sensitive lipids were explicitly modeled with *κ*_app_, the softening effect would be incorrectly double-counted.

## V. DATA AVAILABILITY

All-atom and coarse-grained simulations are available on Zenodo (DOIs: 10.5281/zenodo.10367639 and 10.5281/zenodo.10366938, respectively).

## Supporting information

Supplemental derivations and plots

## ACKNOWLEDGMENTS

This work was supported by the Intramural Research Program of the *Eunice Kennedy Shriver* National nstitute of Child Health and Human Development (NICHD) at the National Institutes of Health.

A.H.B. was supported by a Postdoctoral Research Associate (PRAT) fellowship from the National Institute of General Medical Sciences (NIGMS), award number 1Fi2GM137844-01.

This study utilized the high-performance computational capabilities of the Biowulf Linux cluster at the National Institutes of Health, Bethesda, MD (https://hpc.nih.gov).

