## Supplemental derivations and plots for "Softening in two-component lipid mixtures by spontaneous curvature variance"

### I. Multi-component diffusional softening

Here we derive the energy of density fluctuations and the softening expression for a multi-component system. For both we follow the methodology in the supplementary materials of previous work describing diffusional softening for 2 components [1].

#### A. Density Fluctuations

First we must determine the energy of fluctuations in the density of a multi-component system. We start with 3 components, assuming a total of $N$ lipids in a thin strip of the membrane containing 3 types of lipid with (global) mole fractions $\chi_1$, $\chi_2$, and $\chi_3$, where $\chi_1 + \chi_2 + \chi_3 = 1$, and spontaneous curvature $c_{01}$, $c_{02}$, and $c_{03}$. If $n_1$, $n_2$, and $n_3$ are the number of each type of lipid in the strip, for fixed N we have 2 degrees of freedom, e.g. $n_1$ and $n_2$, with the last value $n_3$ determined by the other two. Assuming $\chi_i$ is the probability of each individual lipid being of type $i$ independent of any other lipids, we find the following probabilities (binomial distribution):

$$\Pr(n_1, n_2) = \Pr(n_1)\Pr(n_2|n_1) \tag{1}$$

$$= \left[\binom{N}{n1}\chi_1^{n_1}(1-\chi_1)^{(N-n_1)}\right]\left[\binom{N-n_1}{n_2}\left(\frac{\chi_2}{1-\chi_1}\right)^{n_2}\left(1-\frac{\chi_2}{1-\chi_1}\right)^{(N-n_1-n_2)}\right] \tag{2}$$

Defining "renormalized" variables for $i = 2$:

$$\tilde{\chi}_2 \equiv \frac{\chi_2}{1-\chi_1} \tag{3}$$

$$\tilde{N}_2 \equiv N - n_1 \tag{4}$$

We can write $\Pr(n2|n1)$ similar to $\Pr(n1)$:

$$\Pr(n2|n1) = \binom{\tilde{N}_2}{n_2}\tilde{\chi}_2^{n_2}(1-\tilde{\chi}_2)^{(\tilde{N}_2-n_2)} \tag{5}$$

The mean and variance of $n_1$ from $\Pr(n1)$ are $\chi_1 N$ and $\chi_1(1-\chi_1)N$, respectively. Similarly, the mean and variance of $n_2$ from $\Pr(n2|n1)$ are $\tilde{\chi}_2\tilde{N}_2$ and $\tilde{\chi}_2(1-\tilde{\chi}_2)\tilde{N}_2$. We approximate the probability of finding $n_1$ and $n_2$ in a bin, in the large N limit, with normal distributions (up to a normalization constant):

$$\Pr(n_1, n_2) \propto \exp(-\frac{n_1 - \chi_1 N}{2\chi_1(1-\chi_1)N})\exp(-\frac{n_2 - \tilde{\chi}_2\tilde{N}_2}{2\tilde{\chi}_2(1-\tilde{\chi}_2)\tilde{N}_2}) \tag{6}$$

For $n_1 \approx \chi_1 N$ the mean and variance of $n_2$ are $\chi_2 N$ and $\chi_2(1-\chi_2)N$, and we can calculate the covariance of $n_1$ and $n_2$:

$$\text{cov}(n_1, n_2) = -\chi_1\chi_2 N \tag{7}$$

In the limit of large $N$ the distribution is well approximated by a bivariate normal distribution

$$\Pr(n1, n2) \propto \exp\left(-\frac{1}{(1-\rho^2)}\left[n_1^2/(2\sigma_1^2) - \rho n_1 n_2/(\sigma_1\sigma_2) + n_2^2/(2\sigma_2^2)\right]\right) \tag{8}$$

with

$$\begin{aligned}\sigma_1 &= \sqrt{\chi_1(1-\chi_1)N} \\ \sigma_2 &= \sqrt{\chi_2(1-\chi_2)N} \\ \rho &= \frac{-\chi_1\chi_2 N}{\sigma_1\sigma_2}\end{aligned} \tag{9}$$

As in Ref. [1], a Fourier representation is chosen for the density inhomogeneity represented by deviations of the components from the background. Finally, instead of representing the energy in terms of $n_1$ and $n_2$, we want the energy in terms of the Fourier amplitudes of the lipid density ($p_{1q}$ and $p_{2q}$). To translate from $n_1$ and $n_2$ into Fourier coefficients $p_{1q}$ and $p_{2q}$ requires transforming from the per-bin variance of $n_1$ and $n_2$, to the variance of the Fourier coefficients of a sampling of those bins (see again Ref. [1]). The ratio of the variances of $p_q$ to $n$ is $\frac{2A}{A_p}$ ($q$-independent), yielding the substitution $n \to \sqrt{\frac{A_p}{2A}}p_q$. The free energy of density fluctuations (for a single sine mode) is:

$$E_\rho = (\frac{k_\mathrm{B}TA_\mathrm{p}}{2A})\Big(1-\frac{\chi_1\chi_2}{(1-\chi_1)(1-\chi_2)}\Big)^{-1}\Big[\frac{p_{1q}^2}{2\chi_1(1-\chi_1)}+\frac{p_{2q}^2}{2\chi_2(1-\chi_2)}+\frac{p_{1q}p_{2q}}{(1-\chi_1)(1-\chi_2)}\Big] \tag{10}$$

#### B. Softening

The total free energy of a single sine mode is:

$$\begin{aligned}E_\mathrm{tot} = \frac{\kappa_\mathrm{int}h_q^2q^4}{2A} + (\frac{\kappa_\mathrm{int}A_\mathrm{p}h_qq^2}{2A})[c_{01}p_{1q}+c_{02}p_{2q}] \\ + (\frac{k_\mathrm{B}TA_\mathrm{p}}{2A})\Big(1-\frac{\chi_1\chi_2}{(1-\chi_1)(1-\chi_2)}\Big)^{-1}\Big[\frac{p_{1q}^2}{2\chi_1(1-\chi_1)}+\frac{p_{2q}^2}{2\chi_2(1-\chi_2)}+\frac{p_{1q}p_{2q}}{(1-\chi_1)(1-\chi_2)}\Big]\end{aligned}$$

where, the first term is the normal Helfrich curvature energy and the second term is the coupling between membrane undulations and density (see the SI of [1]) and the last term is the energetic expression derived above for density fluctuations. We can solve for $p_{1q}$ and $p_{2q}$ that minimize the free energy for a particular value of $h_q$. Substituting those values, we can proceed to calculate $\langle h_q^2\rangle$ and find the softening factor:

$$\alpha = (\frac{\kappa_\mathrm{int}A_\mathrm{p}}{2k_\mathrm{B}T})\Big[c_{01}^2\chi_1(1-\chi_1)+c_{02}^2\chi_2(1-\chi_2)-2c_{01}c_{02}\chi_1\chi_2\Big] \tag{11}$$

Defining mean spontaneous curvature $c_{0\mathrm{m}}$ we can rewrite the softening factor to a form that is invariant under swapping the indices of all the components:

$$c_{0\mathrm{m}} \equiv \chi_1c_{01}+\chi_2c_{02}+\chi_3c_{03} \tag{12}$$

$$\Delta c_{0i} \equiv c_{0i}-c_{0\mathrm{m}} \tag{13}$$

$$\begin{aligned}\alpha = (\frac{\kappa_\mathrm{int}A_\mathrm{p}}{2k_\mathrm{B}T})\Big[\Delta c_{01}^2\chi_1(1-\chi_1)+\Delta c_{02}^2\chi_2(1-\chi_2)+\Delta c_{03}^2\chi_3(1-\chi_3) \\ -2\Delta c_{01}\Delta c_{02}\chi_1\chi_2-2\Delta c_{02}\Delta c_{03}\chi_2\chi_3-2\Delta c_{03}\Delta c_{01}\chi_3\chi_1\Big]\end{aligned}$$

Following a similar line of reasoning this statement can be generalized to the case of n components (assuming similar $A_\mathrm{p}$ for the species):

$$\alpha = (\frac{\kappa_\mathrm{int}A_\mathrm{p}}{2k_\mathrm{B}T})\Big[\sum_{i=1}^{n}\Delta c_{0i}^2\chi_i(1-\chi_i)-2\sum_{i=1}^{n}\sum_{\substack{j=1\\ j\neq i}}^{n}\Delta c_{0i}\Delta c_{0j}\chi_i\chi_j\Big] \tag{14}$$

While the bending modulus is softened, the relative observed curvature is increased by the increased fluctuations:

$$\kappa_\mathrm{app} = \kappa_\mathrm{int}(1-\alpha) \tag{15}$$

$$\langle c\rangle = \frac{A_\mathrm{p}}{A}\Delta c_0(1+\alpha) \tag{16}$$

Inverting Eqs. 15 and 16, to yield $\kappa_{\mathrm{int}}$ and $\Delta c_0$ in terms of $\kappa_{\mathrm{app}}$ and $\langle c\rangle$ is simplified if the relation

$$\langle c\rangle = \frac{A_{\mathrm{p}}}{A}\Delta c_0(1-\alpha)^{-1} \tag{17}$$

is used in lieu of Eq. 16; they are equivalent to first order in $\alpha$. To illustrate how the softening is inverted, consider equations of the form:

$$\begin{aligned} x' &= x(1 - txy_1^2 - \ldots) \\ y_1' &= y_1(1 - txy_1^2 - \ldots)^{-1} \\ &\vdots, \end{aligned}$$

where $x$ and $y$ are substituted to simplify notation. Here $x$ corresponds to a $\kappa$ like term, while the set $y_i$ correspond to curvature terms. The choice of Eq. 17 relates apparent (primed) to intrinsic (unprimed) variables as::

$$xy_i = x'y_i' \tag{18}$$

which is true to first order in $\alpha$ regardless of the assumption. The $\kappa$ form ($x$) is then rearranged as:

$$\begin{aligned} x' &= x(1 - txy_1^2 - \ldots) \\ x' &= x - tx^2y_1^2 - \ldots \\ x' &= x - tx'^2y_1'^2 - \ldots \\ x &= x' + tx'^2y_1'^2 + \ldots \\ x &= x'(1 + tx'y_1'^2 + \ldots) \end{aligned}$$

given

$$\frac{x}{x'} = \frac{y_1'}{y_1} \tag{19}$$

the corresponding relation

$$y_1 = y_1'(1 + tx'y_1'^2 + \ldots)^{-1}$$

holds for curvatures to first order in the coupling such that

$$\kappa_{\mathrm{int}} = \kappa_{\mathrm{app}}(1+\alpha') \tag{20}$$

$$\Delta c_{0,i} = \frac{A}{A_{\mathrm{p}}}\langle c_i\rangle(1+\alpha')^{-1}, \tag{21}$$

also to first order in $\alpha$, with

$$\alpha' = \left(\frac{\kappa_{\mathrm{app}}A^2}{2k_{\mathrm{B}}TA_{\mathrm{p}}}\right)\left[\sum_{i=1}^{n}\langle c\rangle_i^2\chi_i(1-\chi_i) - 2\sum_{i=1}^{n}\sum_{\substack{j=1\\ j\neq i}}^{n}\langle c_i\rangle\langle c_j\rangle\chi_i\chi_j\right] \tag{22}$$

where as opposed to $\alpha$, $\alpha'$ has a squared factor of $\frac{A}{A_{\mathrm{p}}}$ accounting for the proportionality of spontaneous curvature to observed curvature.

The softening expression can be converted to a continuous distribution with fractional density $\chi(c)$ of spontaneous curvature $c$:

$$\alpha = \left(\frac{\kappa_{\mathrm{int}}A_{\mathrm{p}}}{2k_{\mathrm{B}}T}\right)\left[\int c^2\chi(c)\mathrm{d}c - \left(\int c\chi(c)\mathrm{d}c\right)^2\right] \tag{23}$$

where $c_{\mathrm{m}} = \int c\chi(c)\mathrm{d}c$ and the factor $(1-\chi(c))$ is unity when converting from the discrete to continuous case. That is, in the continuous case the fraction of each individual species is vanishingly small compared to unity:

$$\lim_{\Delta\to 0}\sum_{i=-\infty}^{\infty}(\Delta i)^2\chi(i\Delta)\Delta[1-\chi(i\Delta)\Delta] = \int_{-\infty}^{\infty}c^2\chi(c)\mathrm{d}c, \tag{24}$$

for well-behaved $\chi(c)$. It is now clear from Eq. 23 that $\alpha$ depends on the variance of the distribution; this is true in continuous or discrete form. That is, with $\chi(c)$ a distribution with variance $\sigma$,

$$\alpha = \frac{A_{\mathrm{p}}\kappa_{\mathrm{int}}\sigma^2}{2k_{\mathrm{B}}T} \tag{25}$$

### II. Curvature spectrum and number of H-bonds

The curvature spectrum plots for all simulations, showing the q-dependent mean sampled curvature for all lipids of each type and the subgroups with either 0 or 2 H-bonds identified. The histograms of mole fraction of each groups are also shown. (Lipids with 4 or more H-bonds are negligible.)

#### Martini 2

#### 100% DOPC

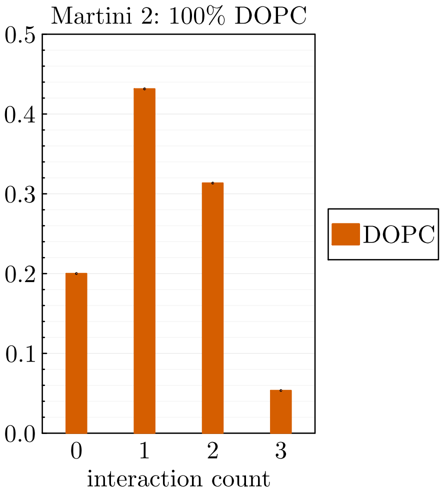


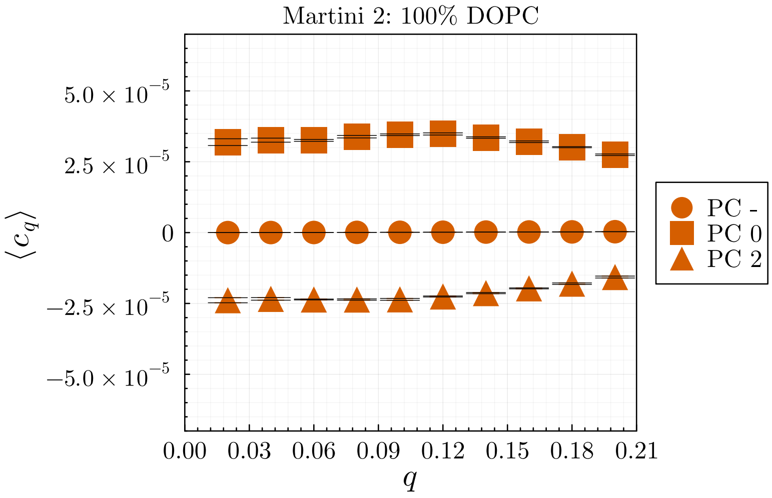


#### 10% DOPE - 90% DOPC

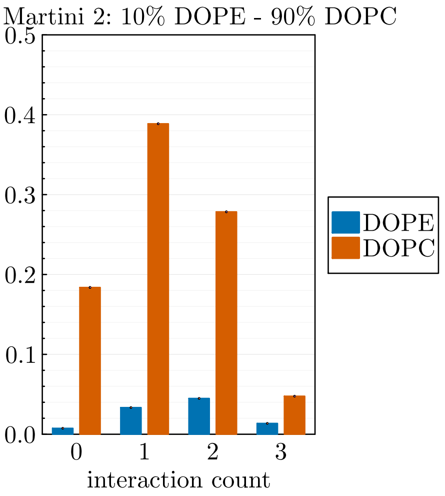


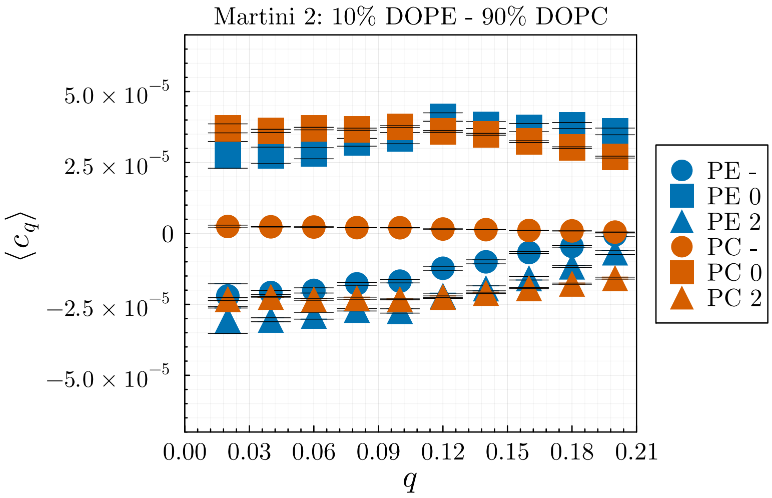

#### 20% DOPE - 80% DOPC

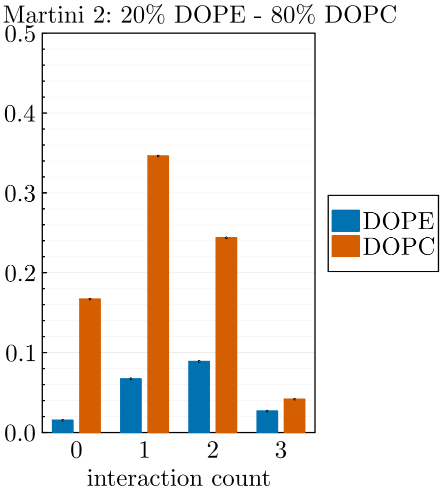


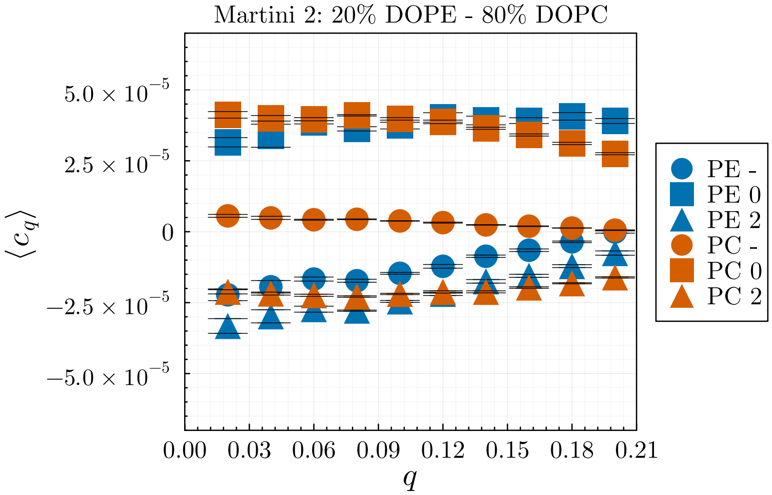


#### 30% DOPE - 70% DOPC

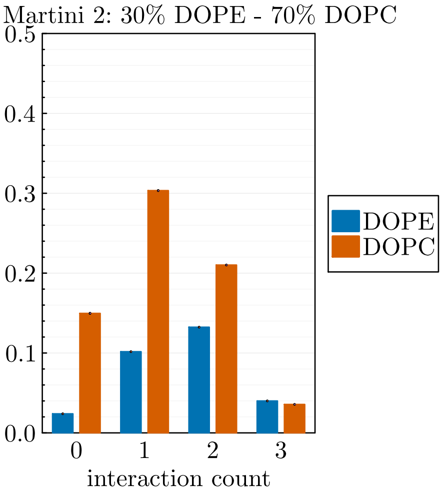


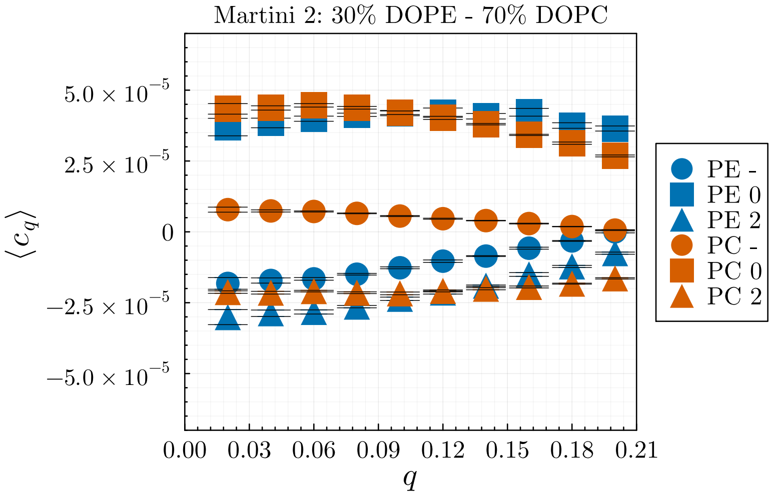


#### 40% DOPE - 60% DOPC

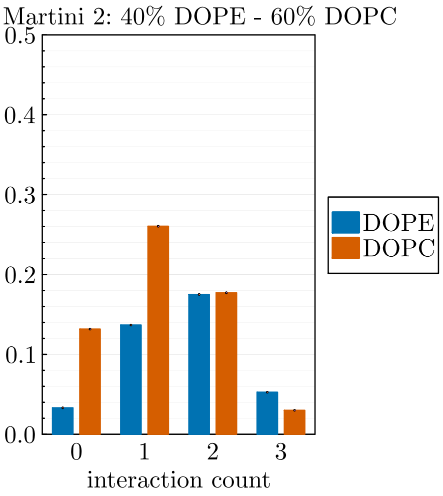


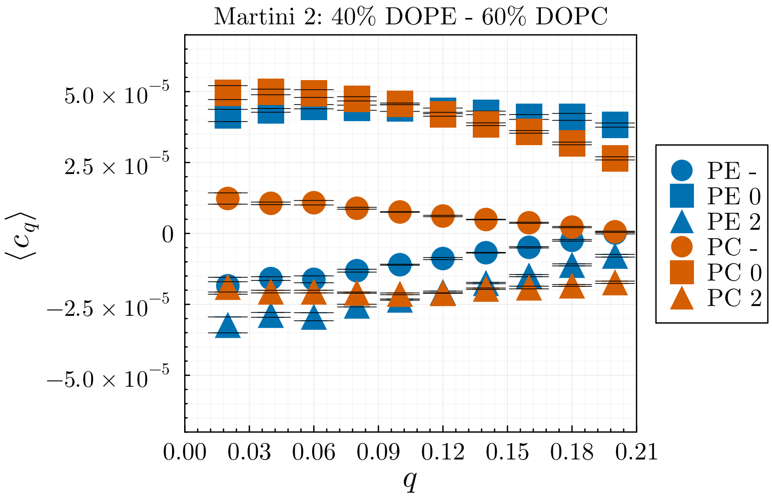

**50% DOPE - 50% DOPC**

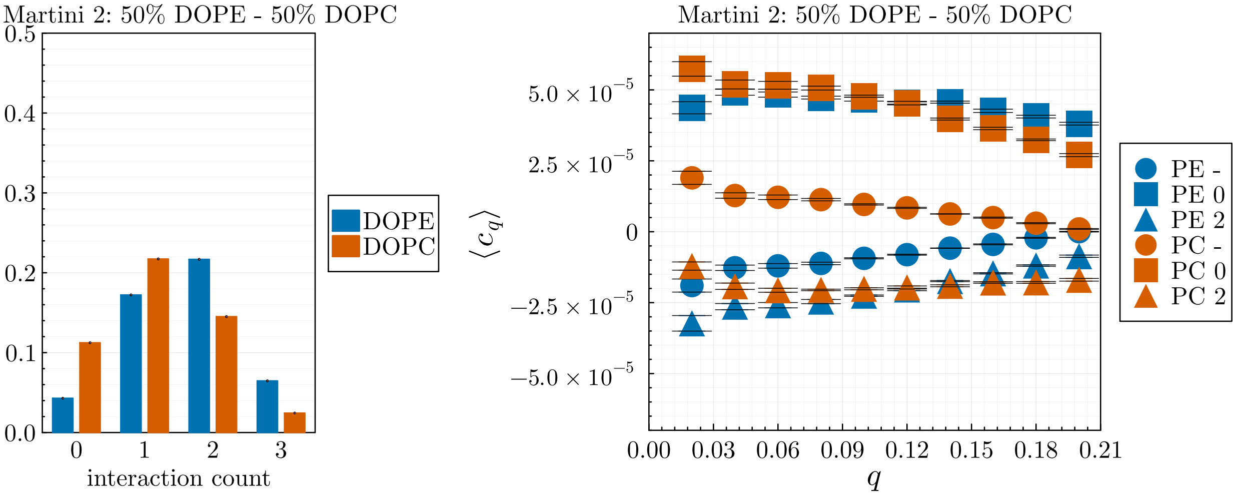


**60% DOPE - 40% DOPC**

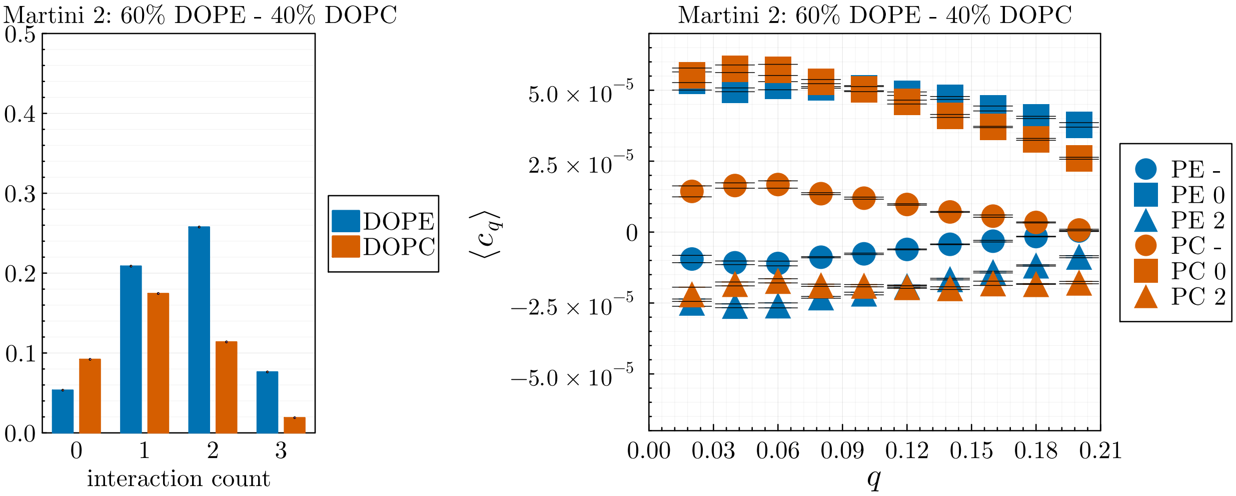


**70% DOPE - 30% DOPC**

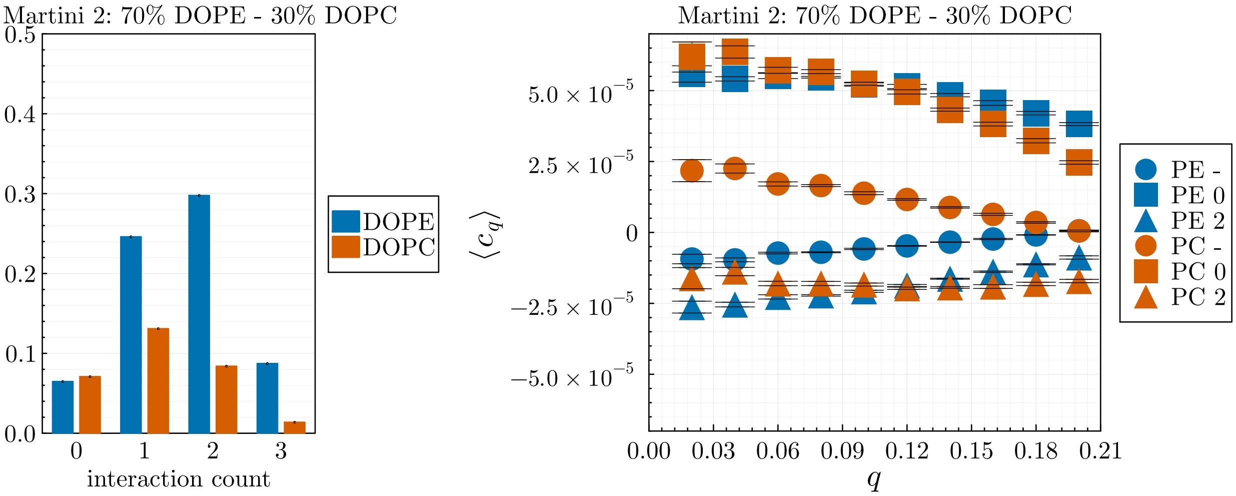

**80% DOPE - 20% DOPC**

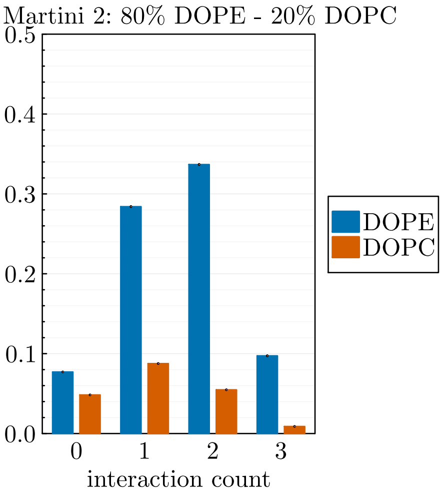


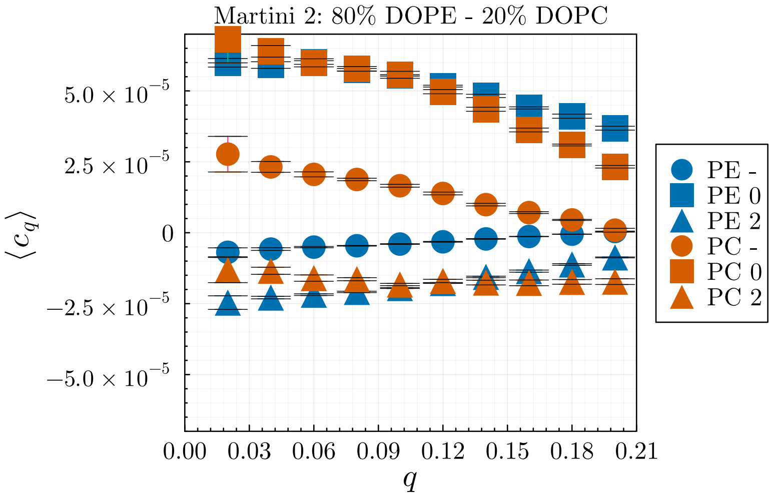


**90% DOPE - 10% DOPC**

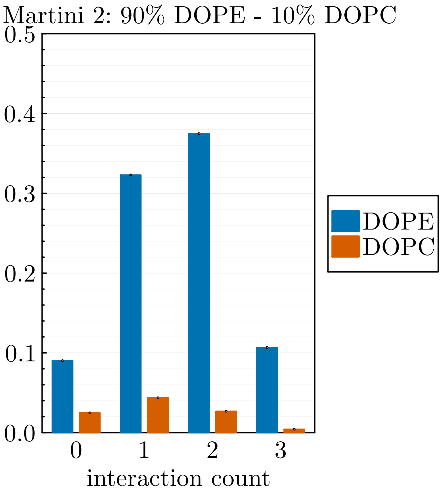


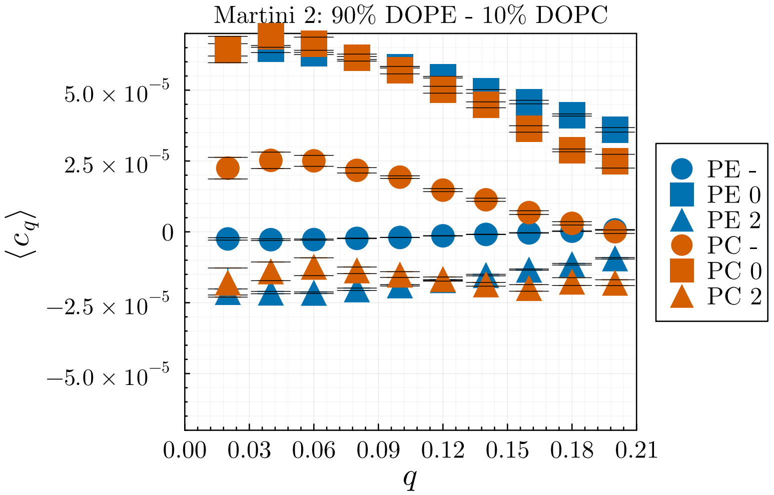


**100% DOPE**

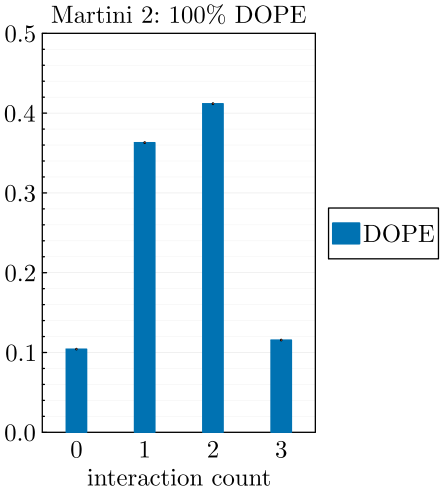


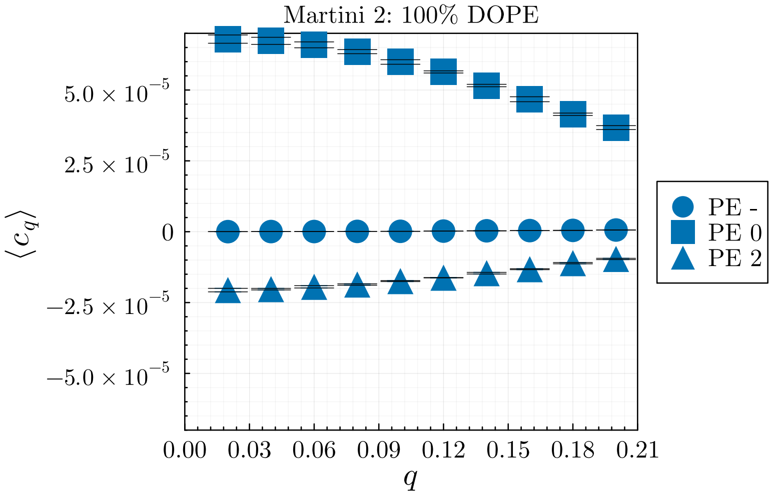

**Martini 3**

### 100% DOPC

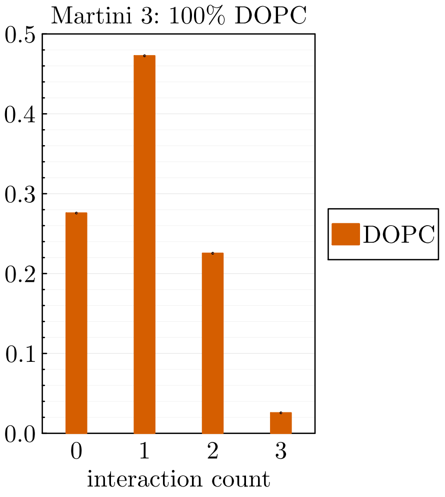


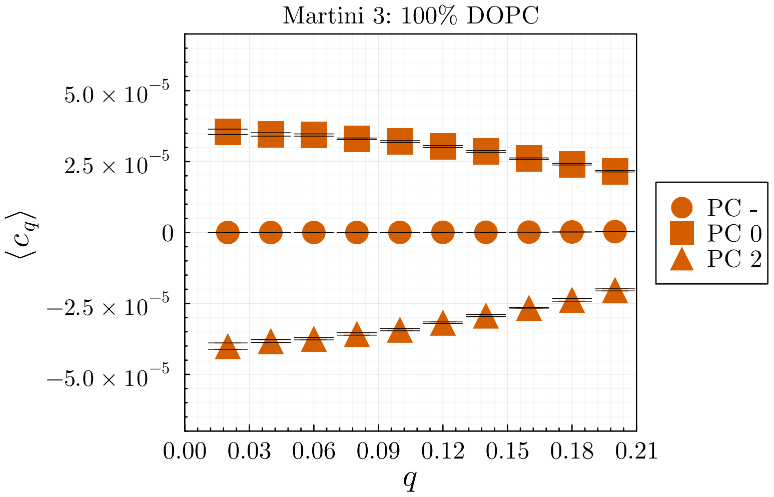


### 10% DOPE - 90% DOPC

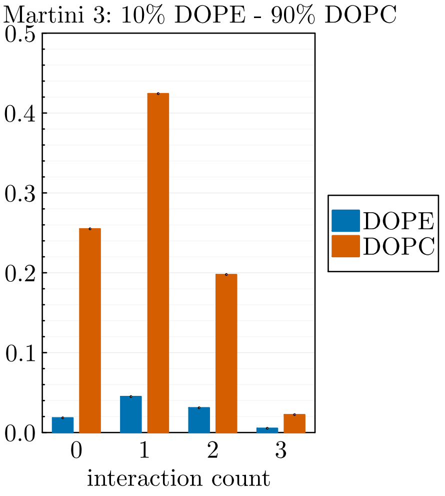


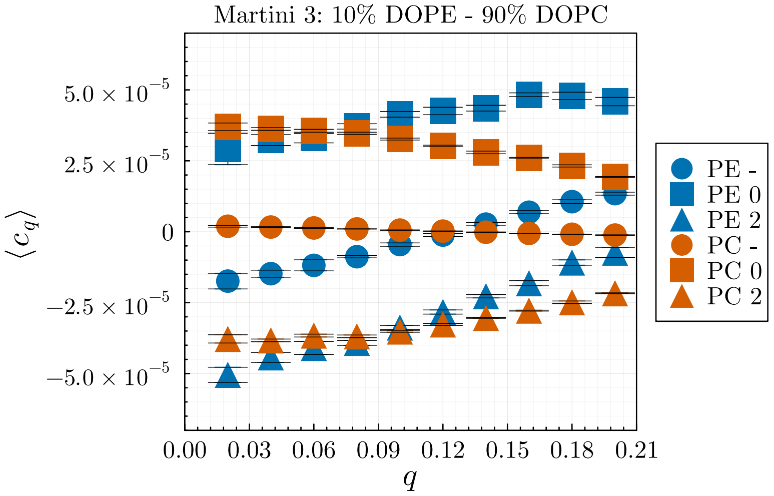


### 20% DOPE - 80% DOPC

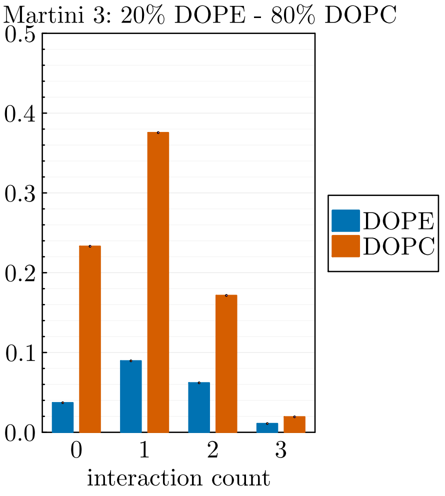


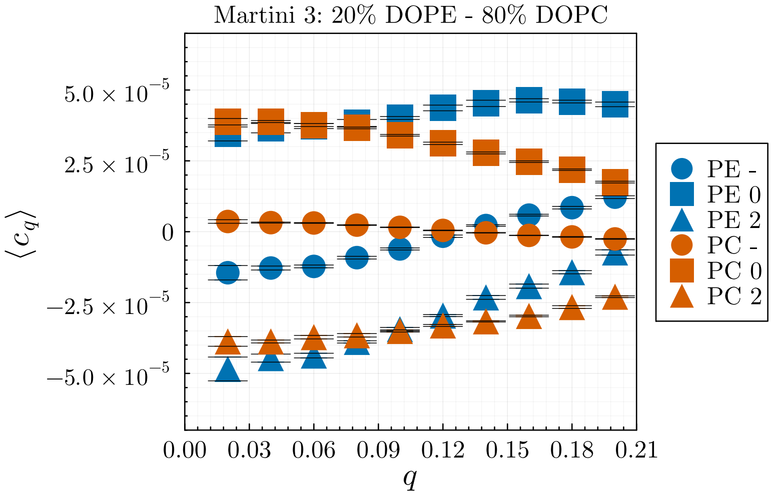

**30% DOPE - 70% DOPC**

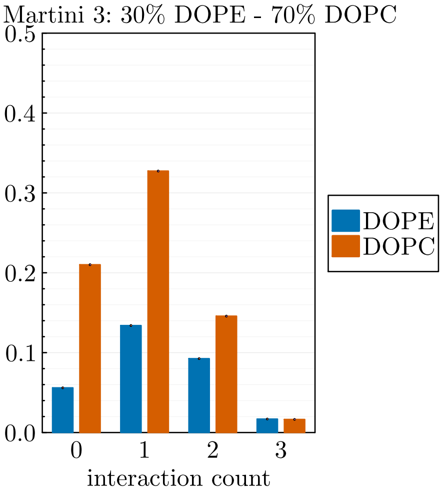


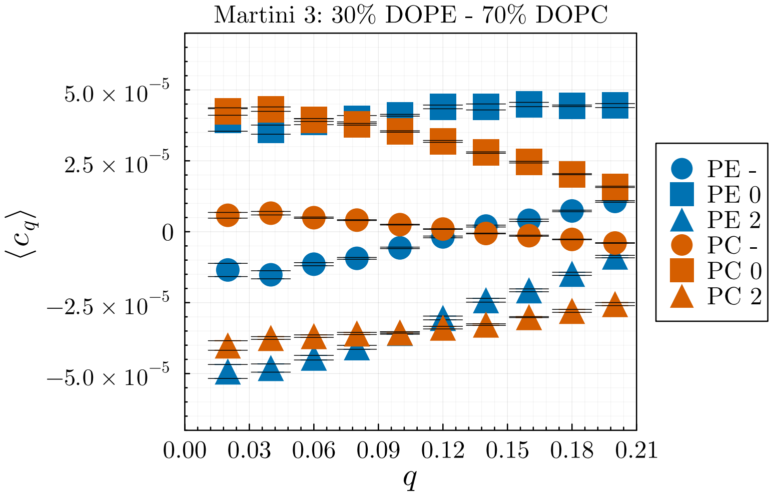


**40% DOPE - 60% DOPC**

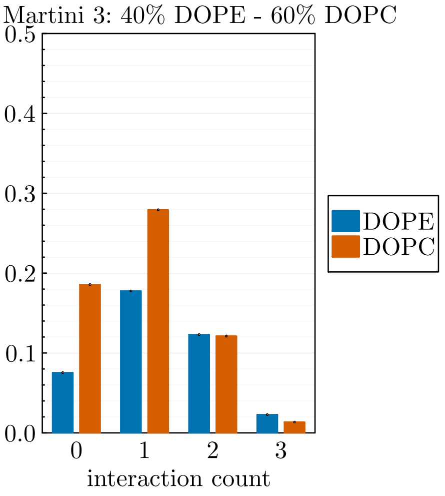


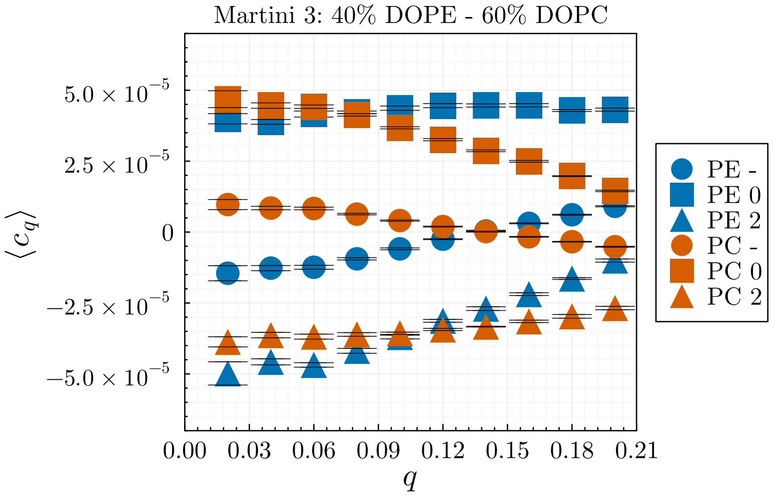


**50% DOPE - 50% DOPC**

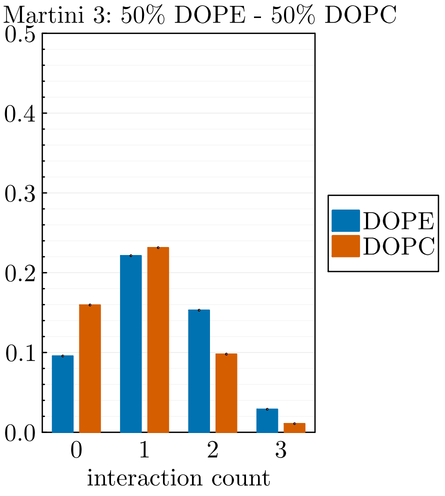


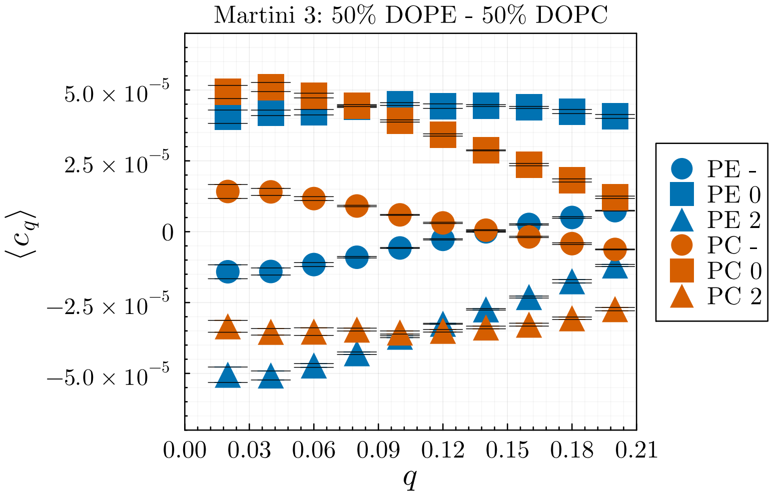

### 60% DOPE - 40% DOPC

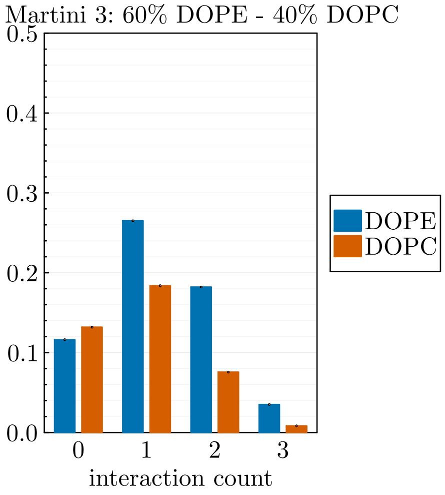


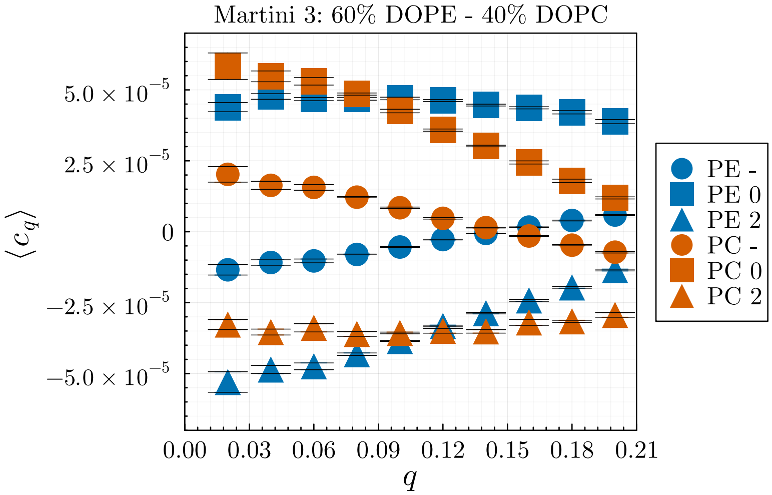


### 70% DOPE - 30% DOPC

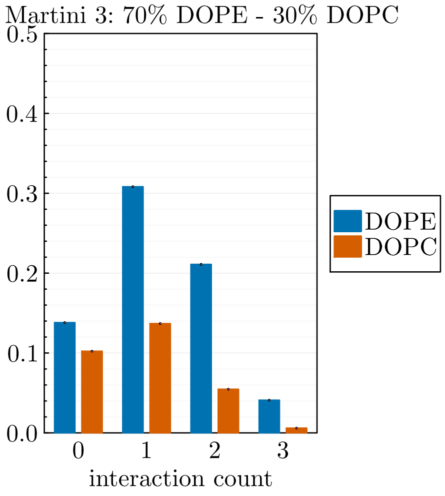


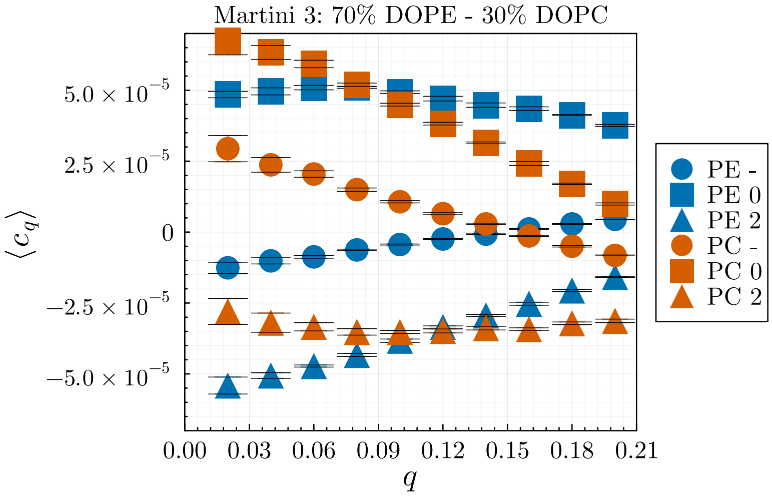


### 80% DOPE - 20% DOPC

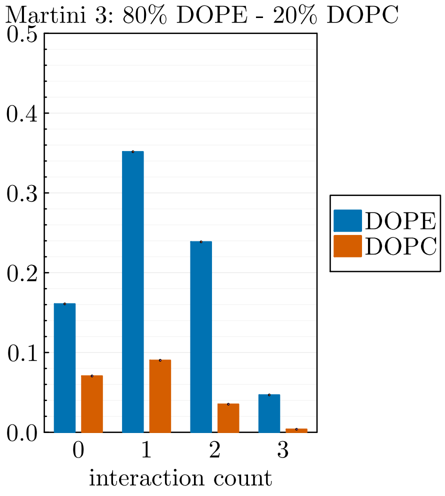


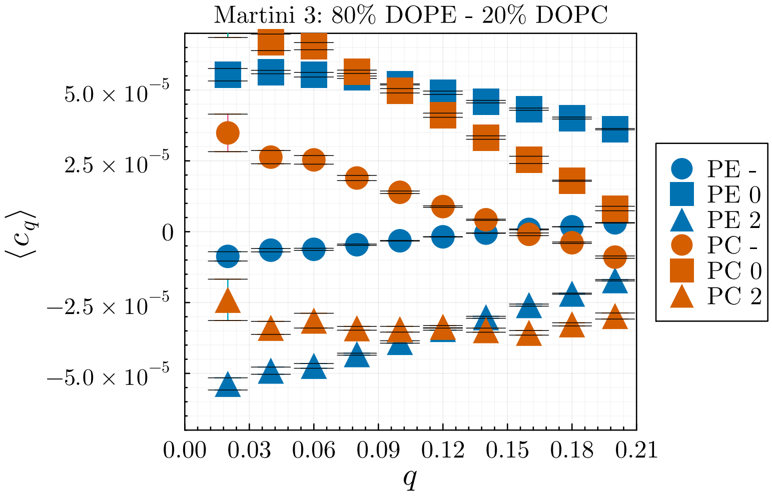

#### 90% DOPE - 10% DOPC

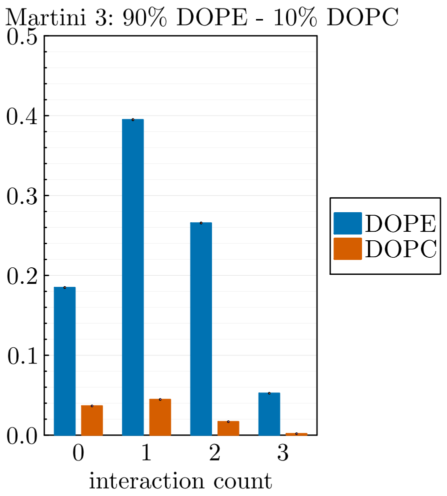


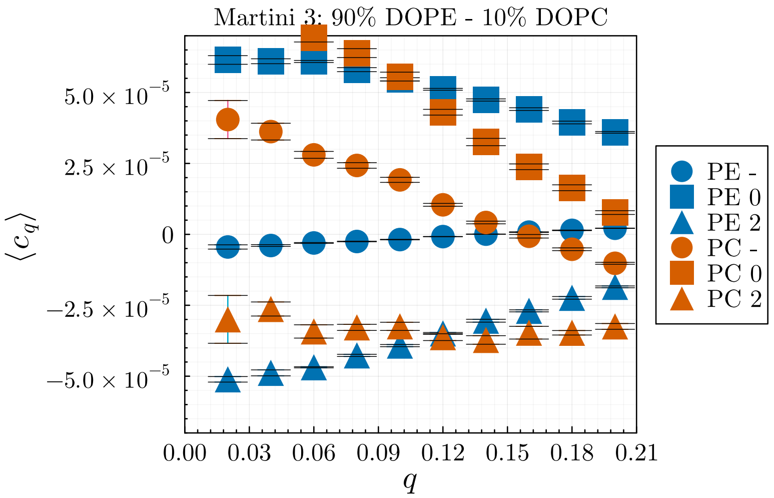


#### 100% DOPE

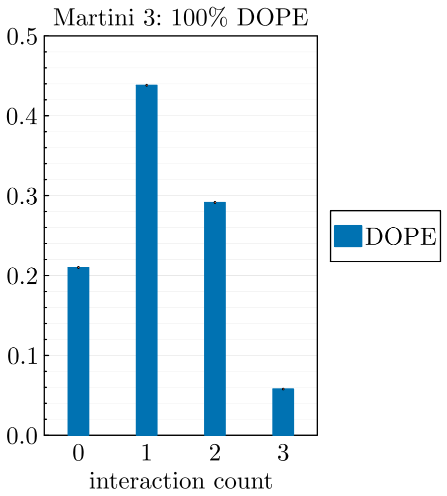


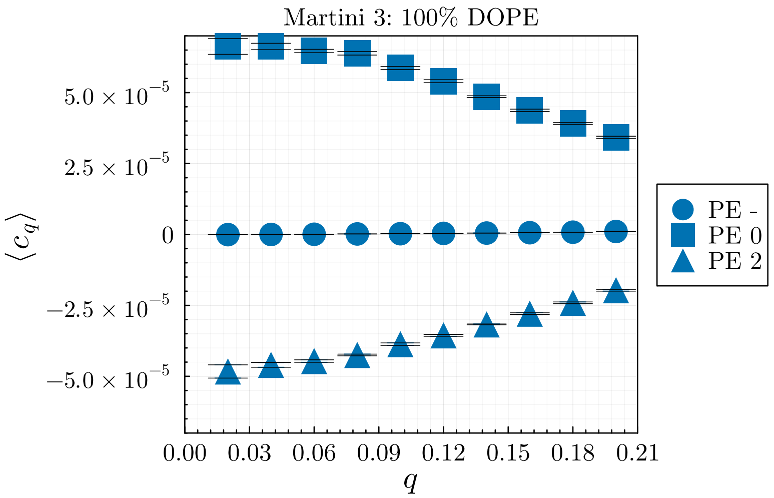


**All-atom**

#### 100% DOPC

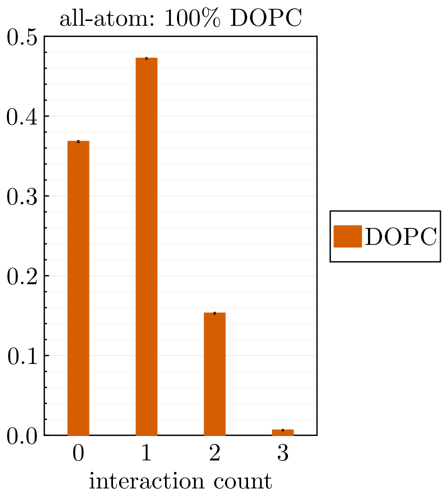


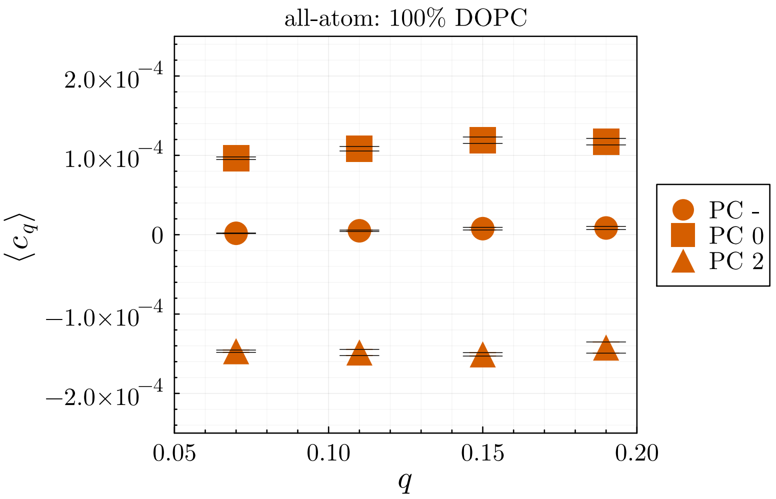

**50% DOPE - 50% DOPC**

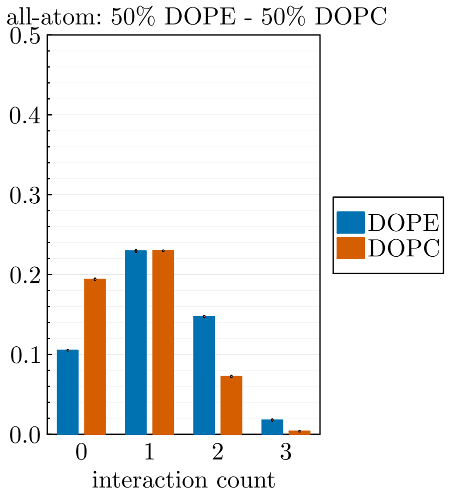


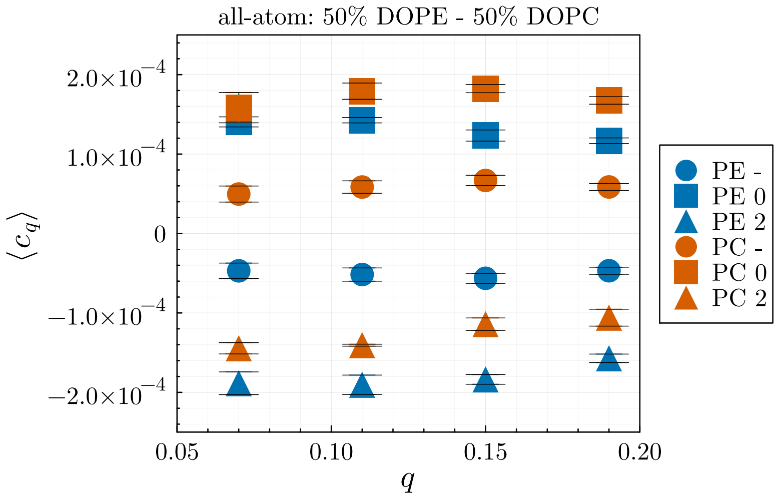


**100% DOPE**

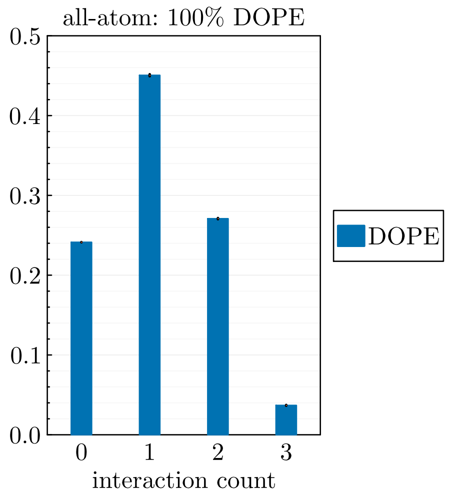


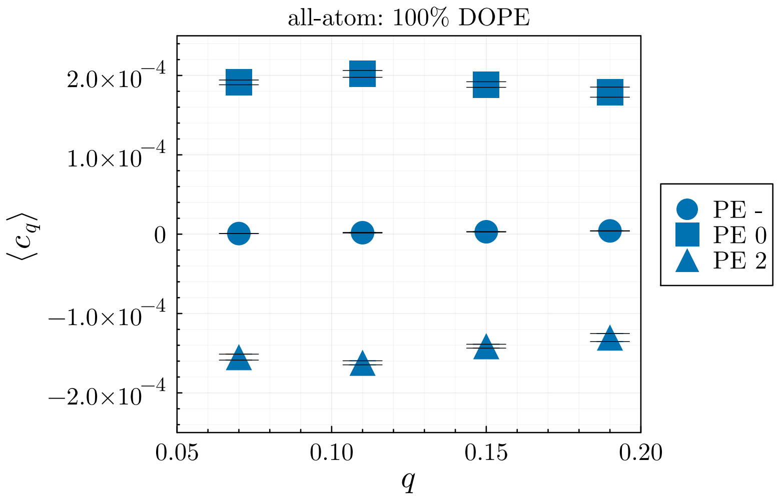


#### III. Bending rigidity from undulation spectrum

We also calculate $\kappa_{\text{app}}$ of the simulated systems (Figs. 1 and 2) using the conventional fluctuation spectrum method of fitting height undulation mode amplitudes to the theoretical prediction. The particular relation we use to extract bending modulus is Eq. 47 from [2], which has corrections for the high $q$ regime.

**Martini**

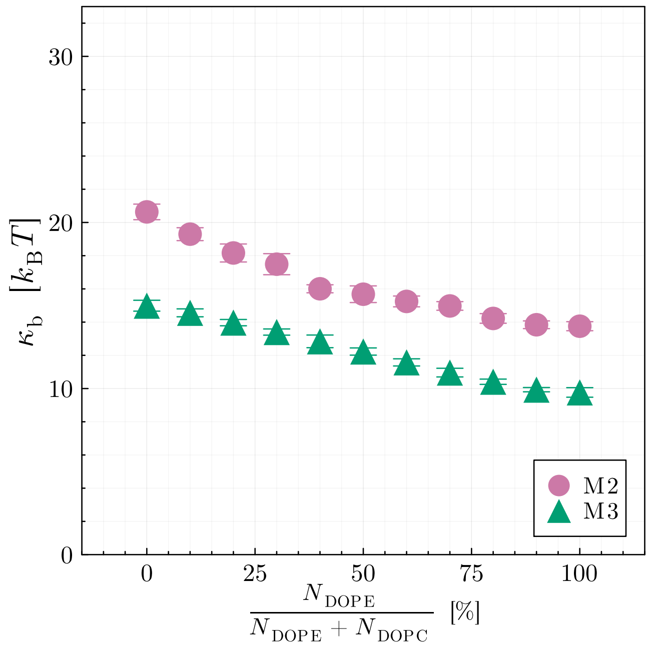


Figure 1: Bending modulus of Martini 2 and 3 simulations computed from the fluctuation method.

**All-atom**

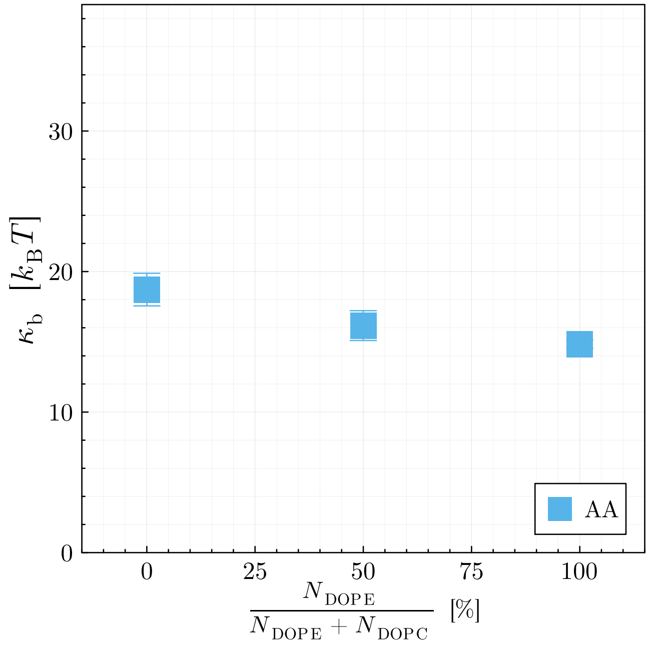


Figure 2: Bending modulus of all-atom simulations computed from the fluctuation method.

[1] H. J. Lessen, K. C. Sapp, A. H. Beaven, R. Ashkar, and A. J. Sodt, Molecular mechanisms of spontaneous curvature and softening in complex lipid bilayer mixtures, Biophysical Journal **121**, 3188 (2022).

[2] M. M. Terzi and M. Deserno, Novel tilt-curvature coupling in lipid membranes, The Journal of chemical physics **147** (2017).
